# Substrate Binding Relieves Autoinhibition of the Mammalian GCN2 Kinase

**DOI:** 10.64898/2026.09.09.750423

**Authors:** Marco A. Catipovic, Rachel Green

## Abstract

Cells respond to amino acid starvation and translational stress through the integrated stress response (ISR) kinase GCN2. Current models maintain that GCN2 senses these stresses either by binding deacylated tRNAs that accumulate during amino acid starvation or by recognizing ribosome collisions that arise from perturbation of translation elongation. As tRNA charging is inherently coupled to translation elongation, assessing the relative contributions of these two pathways has proven difficult. To fully separate the roles of these potential agonists, we reconstituted the regulation of GCN2 *in vitro* using recombinant proteins. GCN2 has low basal activity, which is stimulated by the addition of ribosomes or the ribosomal P-stalk. By contrast, deacylated tRNAs fail to activate GCN2 both *in vitro* and *in vivo* and instead counteract activation by the ribosome and inhibit GCN2 auto- and substrate phosphorylation. This inhibition of autophosphorylation requires the presence of GCN2’s substrate, eIF2α, suggesting that the substrate plays a role in kinase regulation. Indeed, eIF2α binding to a newly defined autoinhibitory loop in the GCN2 kinase domain relieves its repression and stimulates autophosphorylation of the kinase’s activation loop. Together, these data redefine the role of tRNAs in the regulation of GCN2 and reveal a mechanism novel to kinases in general by which substrate binding regulates activation loop phosphorylation.

## Introduction

In metazoans, the four ISR kinases, HRI, PKR, PERK, and GCN2, each respond to distinct cellular stressors by phosphorylating the alpha subunit of translation factor eIF2 (eIF2α), leading to inhibition of translation initiation and induction of the transcriptional integrated stress response (ISR)(*1–3*). GCN2 is activated by amino acid starvation and a variety of ribotoxic stresses that impede translation (*4–8*). Because of the pleiotropic effects of the ISR, GCN2 dysregulation is associated with a variety of cancers and age-related diseases (*9*, *10*); thus, the mechanism of its activation is of considerable interest.

GCN2 is a large dimeric kinase that harbors an N-terminal RWD domain (*11*, *12*) followed by a degenerate pseudokinase domain (PsK) (*13*), an active kinase domain (KD) (*14*, *15*), a tRNA synthetase-like domain (HisRS) (*16–18*), and a ribosome-binding C-terminal domain (CTD) (*19*). Early studies of GCN2 regulation in yeast identified the HisRS domain as necessary for activation and proposed that deacylated tRNAs that accumulated during amino acid starvation could bind this domain to stimulate the kinase (*20–23*). However, more recent studies have demonstrated a tRNA-independent mechanism of GCN2 activation by ribotoxic stresses that stall a fraction of ribosomes leading to collisions between stalled and translating ribosomes on the same mRNA (*6–8*, *24–26*). These collisions recruit GCN2 to the ribosome through its obligate cofactor GCN1 (*26*, *27*), and subsequent interactions with the ribosomal P-stalk proteins activate the kinase (*28–30*).

These models are not mutually exclusive, and separating the contributions of tRNA and ribosomes to this process in cells has proven difficult. Deacylated tRNAs accumulate when cells are starved for amino acids, but the resulting paucity of charged tRNAs also stalls ribosomes and causes collisions (*8*). Thus, it remains unclear which of these two potential agonists is responsible for kinase activity, or whether there is a cooperative role for coincidental tRNA deacylation and ribosome collisions (*24*, *26*, *31*). *In vitro* reconstitution permits independent evaluation of these agonists, but these assays have been fraught. Recombinant yeast GCN2 is constitutively active *in vitro*, preventing assessment of external agonists (*23*, *32*, *33*). Mammalian GCN2 is more amenable to reconstitution, and previous studies have shown tRNA to be a relatively weak agonist compared to ribosomes (*29*, *30*).

However, these reconstitutions monitored eIF2α phosphorylation alone and failed to detect the initial GCN2 autophosphorylations that license kinase activity (*29*, *30*). Activation of GCN2, like most kinases, requires phosphorylation of conserved residues in the kinase domain activation loop, T899 and T904 in human GCN2 (*34*), which induces conformational changes in the kinase domain to make the active site accessible to ATP and eIF2α binding and competent for catalysis (*35*, *36*). While a hallmark of activated GCN2 *in vivo*, previous studies have been unable to detect activation loop phosphorylation of full-length GCN2 *in vitro* and have thus been blind to the regulation of these critical initial events.

Here, using *in vitro* reconstitution with purified components and complementary *in vivo* experiments, we define the contributions of ribosomes and tRNA to the initial activation of GCN2 and subsequent phosphorylation of eIF2α. Surprisingly, we find a crucial role for the substrate itself in the process and thereby describe a novel mechanism of kinase regulation wherein binding of eIF2α to a regulatory loop of GCN2 relieves autoinhibition of the kinase domain.

## Results

### GCN2 Activation by the Ribosomal P-stalk

Previous reconstitutions showed 80S ribosomes to be sufficient agonists of GCN2 *in vitro* (*29*, *30*). Indeed, incubation of recombinant wild-type GCN2 (**Fig S1A**), but not the catalytically dead D848N mutant, with purified ribosomes leads to autophosphorylation of GCN2 as well as phosphorylation of an established model substrate, the N-terminal domain of eIF2α (**Fig S1B, C, D**) (*37*, *38*). In fact, the entirety of the ribosome is dispensable; recombinant P-stalk heteropentamer consisting of uL10, P1, and P2 yields similar activation (**Fig S1B; Fig 1A, B**) (*29*). These activities can be resolved by western blotting with phospho-specific antibodies, autoradiography to detect *^32^*P transferred from γ-phosphate-labeled ATP, or by the retarded migration of the phosphorylated eIF2α in an SDS-PAGE gel (**Fig 1B; Fig S1D**).

**Figure 1.**
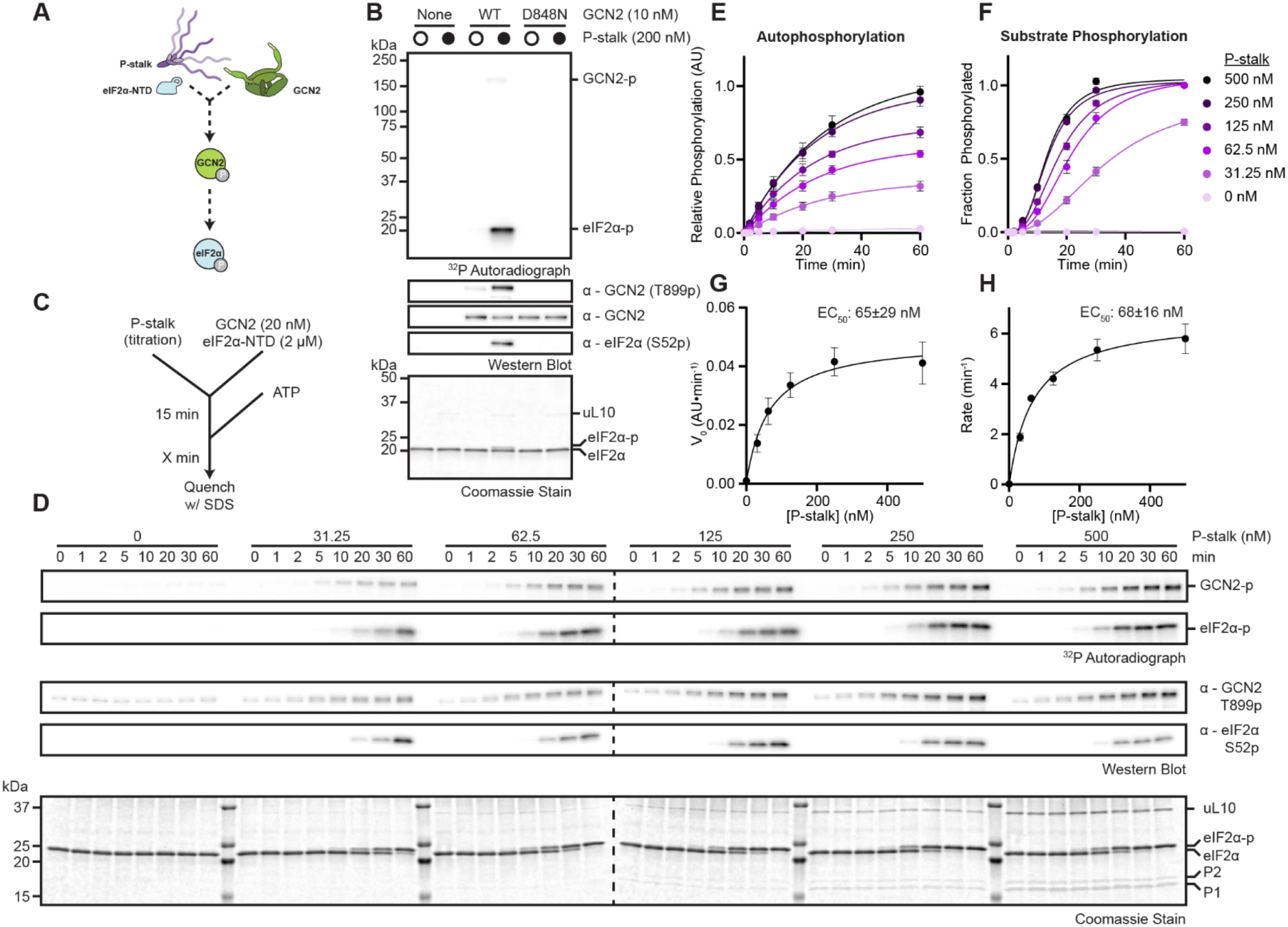
GCN2 Activation by the Ribosomal P-stalk. **(A)** Schematic of a kinase assay using recombinant GCN2 (green), P-stalk (purple), and eIF2α-NTD (light blue). **(B)** Coomassie stain, autoradiograph, and immunoblots with the indicated antibodies of kinase assays as in (A) in the presence or absence of 200 nM P-stalk with 10 nM wild-type (WT) or kinase-dead D848N GCN2 variants and 2 µM eIF2α-NTD. **(C)** Flow chart of kinetic experiment based on kinase assays schematized in (A). **(D)** Coomassie stain, autoradiographs, and immunoblots with the indicated antibodies of kinetic kinase assays with a titration of P-stalk as outlined in (C). **(E)** Intensities of autophosphorylation autoradiographs as in (D) plotted and fit with single exponential functions. Error bars represent S.E.M. of 4 independent replicates. **(F)** As in (E), but with substrate phosphorylation autoradiographs normalized to fraction of phosphorylated eIF2α-NTD visible by Coomassie staining at 60 min. **(G)** V_0_ in (E) as a function of P-stalk concentration fit with a 3-term agonist dose response curve (see Methods). Error bars represent S.E.M. of 4 different replicates. Error on reported EC_50_ is the standard error of the fit. **(H)** As in (G), but with V_Max_ of substrate phosphorylation in (F).

Kinetic analyses of these reactions show both auto- and eIF2α phosphorylation by GCN2 to depend on P-stalk concentration (**Fig 1C, D, E, F**). GCN2 phosphorylation accumulates immediately upon the addition of ATP, whereas substrate phosphorylation exhibits a lag phase characteristic of kinases reliant on autophosphorylation for their activation (**Fig S1E**) (*39*). As previously shown (*34*), and consistent with this dependency, Alanine substitution of either conserved activation loop Threonine abolishes kinase activity (**Fig S1F**). Analysis of the rate of both auto- and substrate phosphorylation yields the P-stalk concentration dependence of each phosphorylation event (**Fig 1G, H**). V_0_ of autophosphorylation can be determined from the slope at the initial timepoints, but due to its initial delay phase, the kinetics of substrate phosphorylation are best fit with two-step hypo-exponential functions, reflective of the sequential nature of the process. The maximum of the derivative of these fits serves as a proxy for V_0_. Both the rates of autophosphorylation and substrate phosphorylation share similar dependencies on P-stalk concentration (**Fig 1G, H**).

### GCN2 Inhibition by tRNA

Deacylated tRNAs, in contrast, do not activate GCN2 *in vitro*. GCN2 has minimal auto- and substrate phosphorylation activity in the absence of P-stalk or ribosomes, visible only by long exposures of autoradiographs (**Fig 2A, B**). Incubation of the kinase with total purified tRNAs deacylated by alkaline hydrolysis fails to produce increased signal for phospho-eIF2α or phospho-GCN2 (**Fig 2A, B**). In fact, tRNAs slightly inhibit the low basal activity of the kinase (**Fig 2B**). This inhibition by tRNA is even more pronounced in the presence of an agonist, such as the P-stalk. Time resolved kinase assays of GCN2 stimulated by multiple P-stalk concentrations in the presence of a tRNA titration show inhibition of both auto- and substrate phosphorylation with increasing concentrations of tRNAs (**Fig 2C, D, E, F, G, H; Fig S2A, B**). This inhibition is not attributable to a contaminant in the tRNA preparation, as *in vitro* transcribed tRNAs also inhibit GCN2 (**Fig S2C**). Nor is it an artifact of kinase assays using only the recombinant N-terminal domain of eIF2α, as phosphorylation of purified endogenous eIF2 trimer is also repressed by tRNA (**Fig S2D**). This regulation is likely specific to GCN2 amongst the ISR kinases since auto- and substrate phosphorylation by the constitutively active fusion of the PERK kinase domain with GST (*40*) is not inhibited by tRNA (**Fig S2E**). Together, these *in vitro* experiments challenge the longstanding notion that deacylated tRNAs activate GCN2.

**Figure 2.**
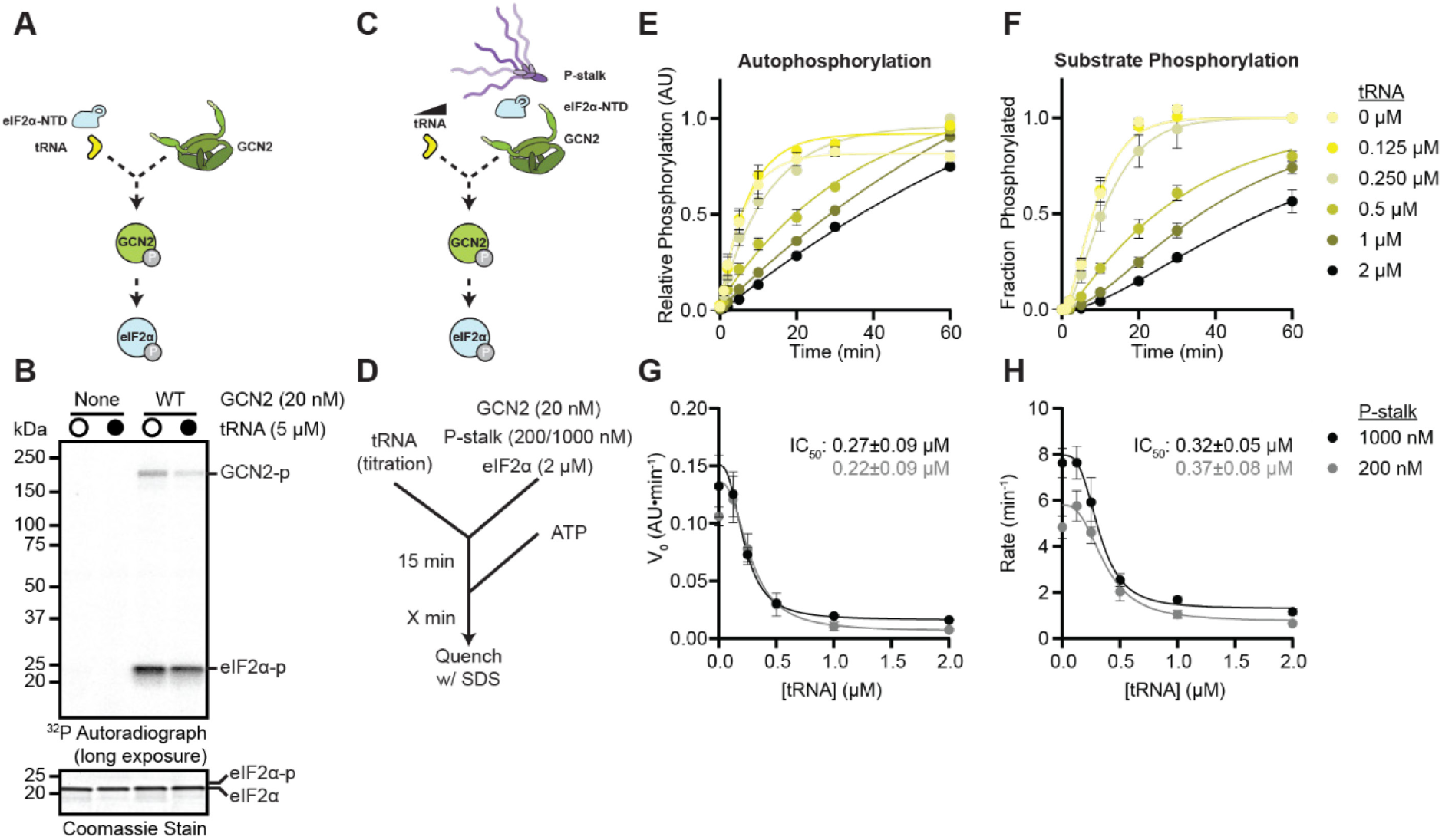
GCN2 Inhibition by tRNA. **(A)** Schematic of a kinase assay using recombinant GCN2 (green), eIF2α-NTD (light blue), and purified deacylated pig liver tRNA (yellow). **(B)** Coomassie stain and autoradiograph (72 h exposure) of kinase assays as in (A) in the presence or absence of tRNA. **(C)** Schematic of a kinase assay as in (A) with the addition of P-stalk (purple). **(D)** Flow chart of kinetic experiment based on kinase assays schematized in (C). **(E)** Intensities of autophosphorylation autoradiographs from experiments as in (D) with 1000 nM P-stalk (Figure S2A) plotted and fit with single exponential functions. Error bars represent S.E.M. of 2 independent replicates. **(F)** As in (E) but with substrate phosphorylation autoradiographs normalized to fraction of phosphorylated eIF2α-NTD visible by Coomassie staining at 60 min. **(G)** V_0_ in (E) as a function of P-stalk concentration fit with a 4-term inhibitor dose response curve (see Methods). Error bars represent S.E.M. of 2 different replicates. Error on reported IC_50_ is the standard error of the fit. **(H)** As in (G), but with V_Max_ of substrate phosphorylation in (F).

The dependence of translation elongation on charged tRNA levels in cells complicates testing the direct activation or inactivation of GCN2 by deacylated tRNA *in vivo*. While cellular treatments such as addition of translation elongation inhibitors or depletion of release factors likely stall ribosomes without perturbing tRNA charging, deacylating tRNAs without affecting translation is more challenging. Amino acid starvation or tRNA synthetase inhibitors lead to deacylation of specific tRNAs, but they also stall ribosomes at codons decoded by the missing amino-acyl tRNAs (*8*, *41*, *42*). Bacterial amber suppression tRNAs engineered to be bio-orthogonal to mammalian tRNA synthetases provide a mechanism to isolate these processes. These tRNAs are only charged by cognate bacterial synthetases in the presence of specific, frequently non-canonical, amino acid analogs (**Fig 3A**) (*43–45*). As GCN2 shows little specificity in its tRNA binding (*23*), introduction of these tRNAs without their cognate synthetase or amino acid should activate the kinase if deacylated tRNAs alone are sufficient agonists.

**Figure 3.**
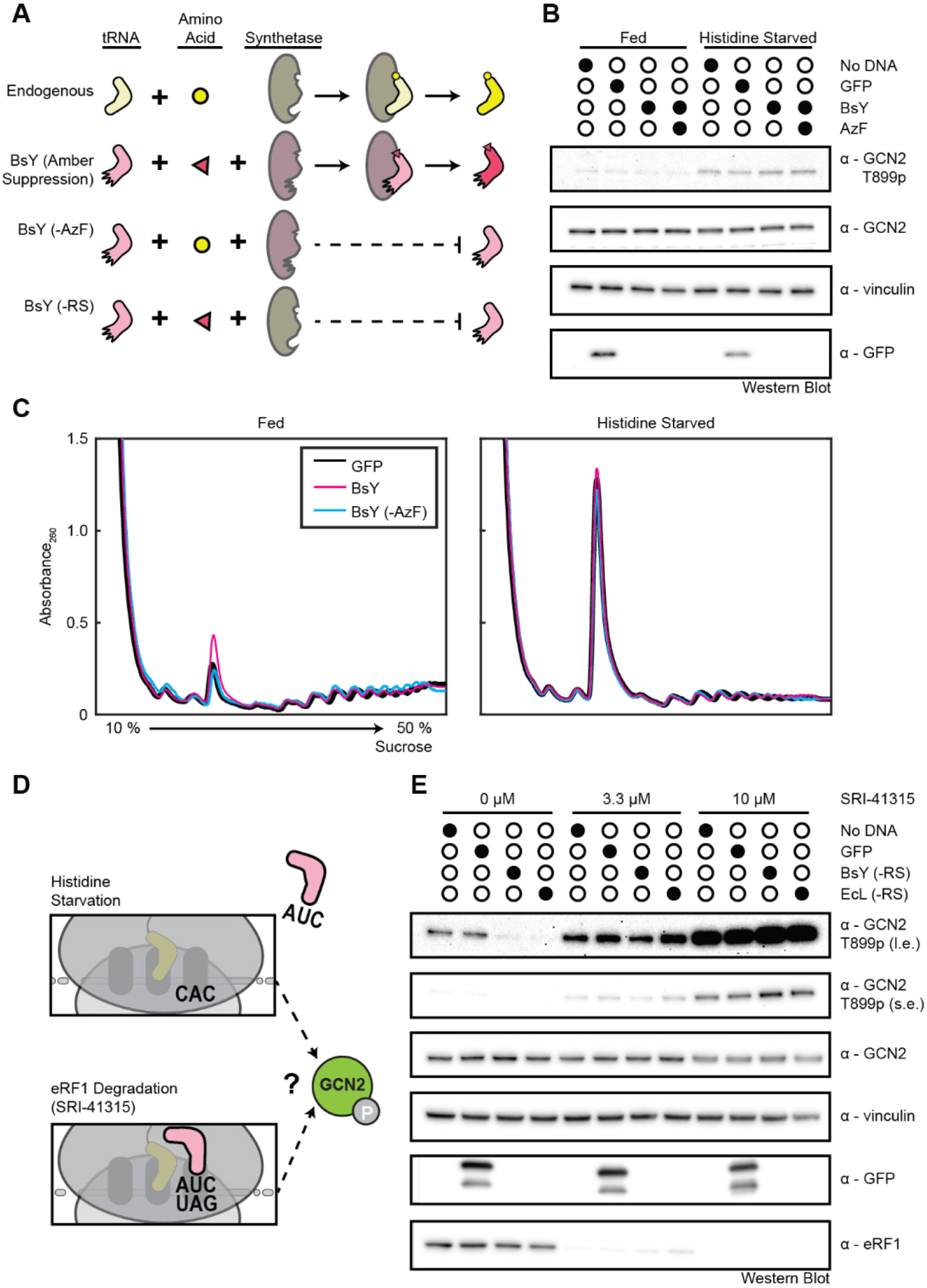
Deacylated tRNAs do not Activate GCN2 *In vivo*. **(A)** Charging of endogenous and amber suppressor tRNAs in cells. Amber suppressor tRNAs (BsY) are only charged in the presence of their cognate synthetase (RS) and their intended amino acid (AzF). **(B)** Immunoblots of GCN2 activity in ΔPKR HEK293T cells transfected with no DNA, GFP (pFD042), or BsY amber suppression plasmids (pIRE4-Azi) in the presence or absence of Azido-Phenylalanine (AzF) grown on complete media or starved of Histidine for 1 h. **(C)** Sucrose gradients from ΔPKR HEK293T transfected as in (B) in the presence or absence of AzF grown on complete media or starved of Histidine for 1 h. **(D)** Histidine starvation stalls ribosomes at codons non-cognate to amber suppressor tRNA anticodons. The eRF1 degrader SRI-41315 stalls a fraction of ribosomes at UAG stop codons cognate to amber suppressor anticodons. **(E)** Immunoblots of GCN2 activity in ΔPKR HEK293T cells transfected with no DNA, GFP (pFD042), BsY amber suppression plasmid lacking the RS (pMC389), or EcL amber suppression plasmid lacking the RS (pMC390) and treated with the indicated dosage of SRI-41315 for 4 h to induce eRF1 degradation.

However, *in vitro*, these tRNAs inhibit GCN2 activity as well as endogenous tRNAs (**Fig S3A**). And, while a *Bacillus stearothermophilus* tRNA_Tyr_ (BstY) accumulates in cells when expressed along with its cognate synthetase (RS) in the presence or absence of its bespoke amino acid analog Azido-phenylalanine (AzF) (**Fig S3B**), it fails to activate GCN2 *in vivo*. GCN2 phosphorylation is not enhanced by transfection of plasmids encoding BstY (**Fig 3B**), while a canonical GCN2 trigger, starvation for Histidine (His), readily induces GCN2 autophosphorylation (**Fig 3B**). Furthermore, while activation of the ISR by His starvation inhibits translation initiation leading to accumulation of monosomal ribosomes visible by polysome profiling, expression of BstY does not perturb global translation (**Fig 3C**).

Several studies have proposed that tRNAs provide additional stimulation to GCN2 activated primarily by interactions with the ribosome, rather than serve as independent agonists (*26*, *46*). However, starvation of cells expressing BstY, in the presence or absence of AzF, produces similar levels of phospho-GCN2 and translation inhibition as starvation of control transfected cells (**Fig 3B, C**), suggesting that BstY does not synergize with His starvation to induce more potent GCN2 activation.

Most existing models, though, stipulate that the anticodon sequence of the stimulatory deacylated tRNAs must be cognate to the mRNA codon in the ribosomal A site (*26*, *46*). Histidine starvation stalls ribosomes at CAC and CAU codons, clearly not cognate matches for the amber suppressor tRNAs’ CUA anticodon. A better experiment, then, would test the synergistic activation of GCN2 by these tRNAs and ribosomes stalled at UAG amber stop codons (**Fig 3D**). The small molecule SRI-41315 serves as a molecular glue that triggers the degradation of eRF1, thus trapping ribosomes at stop codons (*47*, *48*). The resulting stalled ribosomes collisions induce GCN2 phosphorylation within 4 hours of drug treatment (**Fig S3C, D**), but transfection of deacylated amber suppressor tRNAs does not modulate this response (**Fig 3E**). Treatment of cells expressing BstY or an alternative amber suppressor tRNA derived from an *Escherichia coli* tRNA_Leu_ (EcL) with low or high dosages of SRI-41315 produces similar levels of phosphorylated GCN2 as observed in un- or GFP transfected control cells (**Fig 3E**). In fact, long exposure of the western blots from control treated cells shows that these exogenously expressed tRNAs inhibit the low basal activity of GCN2, consistent with *in vitro* results (**Fig 3E**, **Fig 2B**).

### Mechanism of GCN2 Inhibition by tRNA

Multiple studies have reported that tRNAs bind GCN2 at the tRNA synthetase-homologous HisRS domain, coordinated by the motif 2 (m2) residues conserved from class II tRNA synthetases (*18*, *20*, *21*, *23*, *29*). Surprisingly, the m2 motif is also required for ribosome binding and activation by the ribosomal P-stalk (**Fig S4A**) (*29*). Thus, a straightforward hypothesis for GCN2 activation by P-stalk and inhibition by tRNAs is that the two factors compete for this binding site. If tRNA binding is inert and the P-stalk stimulatory, then displacement of bound P-stalk by tRNA would manifest as kinase inhibition. However, several lines of evidence contradict this model. Firstly, while Alphafold3 predicts the binding of the acidic tails of P-stalk proteins to the GCN2 m2 motif and successfully places a tRNA_His_ in the proper position on Histidine tRNA synthetase, it struggles to canonically orient a tRNA_His_ bound to the GCN2 HisRS domain (**Fig S4B**). Secondly, if direct competition between these factors were regulating the kinase, then the IC_50_ of tRNA would vary with the P-stalk concentration. However, kinase assays in the presence of a five**-**fold range of P-stalk concentrations show little difference in the inhibitory concentration of tRNA (**Fig 2G, H**). Lastly, if tRNAs and the P-stalk compete for m2 motif binding, then challenging GCN2-bound P-stalk with tRNAs should elute the P-stalk from GCN2, but a titration of tRNA fails to displace P-stalk from Strep-tagged GCN2 immobilized on beads (**Fig S4C**).

Rather, the inhibition of basal kinase activity suggests that tRNAs are inhibitory independent of the P-stalk. This intrinsic inhibition is more evident with GCN2 mutants that are active even in the absence of ribosomes or the ribosomal P-stalk. Genetic suppressor screens in yeast for GCN2 mutants that maintained activity in an m2 mutant background identified two mutations in the kinase domain, R794G and F842L, that rendered the kinase constitutively active (*49*). The corresponding variants of the human protein, S808G and F855L, are active *in vitro* in the absence of P-stalk or ribosomes (**Fig S4D**). However, both mutants are inhibited by tRNA with a concentration dependence comparable to that of the wild-type kinase (**Fig S4E, 2H**). Consistent with its initial identification as an m2 suppressor mutant, the S808G variant functions *in vitro* even when combined with m2 mutations in the HisRS domain, though its activity is dampened compared to the variant with S808G alone (**Fig S4F**). Importantly, despite disruption of the presumed tRNA binding site, this double mutant is still inhibited by tRNA (**Fig S4G**). Overall, GCN2 inhibition by tRNA appears agnostic to interactions involving its m2 motif, which has long been considered a crucial determinant of tRNA binding. This raises the possibility that GCN2 may be inhibited by a broader range of polyanions beyond deacylated tRNA alone. Indeed, GCN2 can be inhibited *in vitro* with predominantly acylated tRNAs (**Fig S4H**), RNAs transcribed *in vitro* from scrambled tDNA sequences (**Fig S4I**), and even the ss- and dsRNA analogs polyA and polyIC (**Fig S4J**).

### Activation of GCN2 by eIF2α

A general model of kinase function dictates that activation of the kinase domain precedes substrate binding (*50*). As such, tRNAs’ inhibition of substrate phosphorylation could simply be a manifestation of their inhibition of autophosphorylation during the initial steps of kinase activation. To address this possibility, we simplified our system and focused on the autophosphorylation of GCN2 and its inhibition by tRNA, leaving out the substrate which seemed unlikely to contribute to these early steps. Strikingly, though, tRNAs added to GCN2 and P-stalk in the absence of eIF2α fail to inhibit the autophosphorylation of GCN2 (**Fig 4A, B, C, D; Fig S5A**), despite being a potent repressor of this step when the substrate is present (**Fig 4E**, **2H**). These observations raise the unprecedented possibility that GCN2 activation requires interactions with eIf2α.

**Figure 4.**
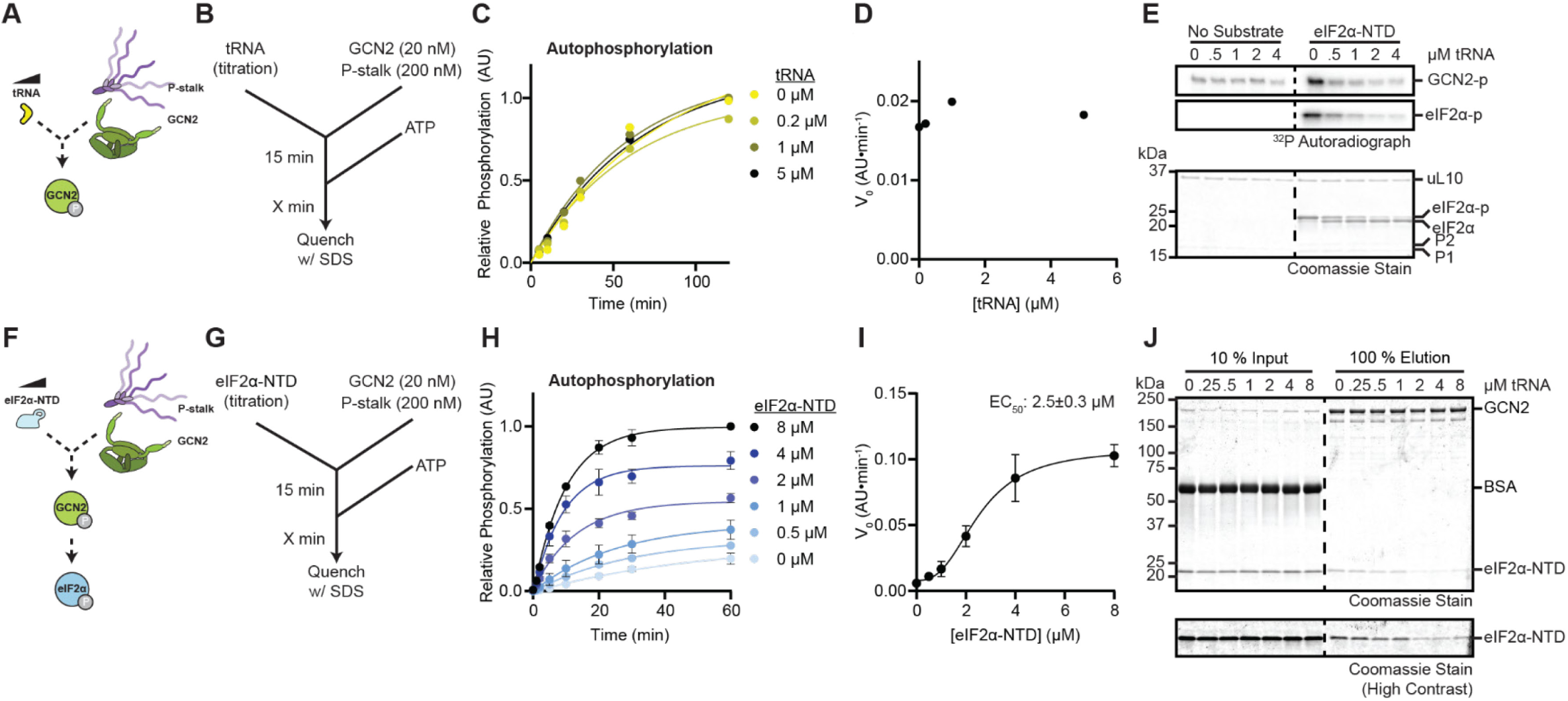
Activation of GCN2 by eIF2α. **(A)** Schematic of kinase assay using recombinant GCN2 (green), P-stalk (purple), and purified deacylated pig liver tRNA (yellow). **(B)** Flow chart of kinetic experiment based on kinase assays schematized in (A). **(C)** Intensities of autophosphorylation autoradiographs from experiments as in (B) with 200 nM P-stalk (Fig S5A) plotted and fit with single exponential functions. **(D)** V_0_ in (C) as a function of tRNA concentration. **(E)** Coomassie stain and autoradiographs from kinase assays as schematized in (A) run for 30 min with 20 nM GCN2 with a titration of tRNA in the presence or absence of 2 µM eIF2α-NTD. **(F)** Schematic of a kinase assay as in (A) but with a titration of eIF2α-NTD (light blue) in place of tRNA. **(G)** Flow chart of kinetic experiment based on kinase assays schematized in (F). **(H)** Intensities of autophosphorylation autoradiographs from experiments as in (G) with 1000 nM P-stalk (Figure S5B) plotted and fit with single exponential functions. Error bars represent S.E.M. of 2 independent replicates. **(I)** V_0_ in (H) as a function of eIF2α-NTD concentration fit with a 4-term agonist dose response curve (see Methods). Error bars represent S.E.M. of 2 different replicates. Error on reported EC_50_ is the standard error of the fit. **(J)** Coomassie stains from co-immunoprecipitation experiments with 200 nM Strep-tagged GCN2 and 2 µM eI2Fα-NTD in the presence of a titration of tRNA. The region including eIF2α-NTD is displayed at two separate contrast levels for clarity.

If GCN2 is indeed activated by its substrate, then its autophosphorylation should be sensitive to the concentration of eIF2α. Time resolved kinase assays with a titration of eIF2α directly test this proposition (**Fig 4F, G**; **Fig S5B**). As anticipated, the efficiency of substrate phosphorylation depends on the concentration of substrate, and accordingly, these experiments provide an estimate of GCN2’s K_m_ for eIF2α-NTD (**Fig S5C, D**). However, the rate of GCN2 autophosphorylation also scales significantly with the concentration of eIF2α-NTD or purified endogenous eIF2 trimer (**Fig 4H; S5B, E**), with an EC_50_ for eIF2α-NTD near 2 µM (**Fig 4I**). These data are consistent with the substrate potentiating the kinase’s activation.

Given this role for eIF2α in activating GCN2, it is possible that the observed inhibition by RNA could result from RNA displacing bound substrate from the kinase. GCN2 and eIF2α co-immunoprecipitation experiments test this hypothesis. Indeed, eIF2α binds GCN2 *in vitro*, even in the absence of P-stalk or ATP (**Fig S5F**), and competition with a titration of tRNA partially displaces the bound eIF2α-NTD (**Fig 4J**). Together, these data suggest that eIF2α is an activator of GCN2 and that RNAs inhibit GCN2 activation by disrupting a regulatory interaction necessary for autophosphorylation of the kinase.

### GCN2 Determinants of Activation by Substrate

Based on its homology to other ISR kinases, GCN2 likely binds eIF2α through the C-terminal lobe of its kinase domain to catalyze its phosphorylation. Crystal structures of the related kinase PKR bound to eIF2α-NTD, as well as Alphafold3 predictions of the GCN2 KD-substrate complex, both show the S1 domain of eIF2α cradling the αG helix of the kinase domains (**Fig S6A**) (*38*). Intriguingly, in some crystal structures of the substrate-free GCN2 kinase domain, the dephosphorylated activation loop (yellow) lies over the αG helix in a position that would clash with substrate binding (**Fig S6A**) (*15*). These interactions between its activation loop and αG helix could maintain GCN2 in an inhibited conformation. In this model, kinase activation would be facilitated by substrate binding which displaces this loop, freeing it for autophosphorylation, and activating the kinase.

The activation loop of GCN2 is significantly longer than those of the other ISR kinases, especially PKR (**Fig S6B**), which renders it uniquely able to engage the αG helix. If this interaction is indeed repressive, chimeric GCN2 constructs with the endogenous activation loop substituted with those of HRI, PKR, or PERK should be primed for autophosphorylation independent of substrate binding and thus should see no stimulation in the presence of eIF2α. However, while these constructs have overall reduced activity relative to wild-type GCN2, autophosphorylation of each of these chimeras is still accelerated by eIF2α (**Fig S6C**), suggesting that the interaction between the activation loop and the αG helix is not responsible for substrate-mediated stimulation of GCN2 autophosphorylation.

Furthermore, a previously described mutation of Thr924 within the αG helix of yeast GCN2 impairs substrate phosphorylation without perturbing autophosphorylation (*51*). Mutation of the corresponding Thr943 in human GCN2 to Ala or Asp also disrupts substrate phosphorylation, but the autophosphorylation of these variants is only slightly reduced and still depends on the substrate (**Fig S6D**). As autophosphorylation of kinase mutants with an αG helix incapable of coordinating substrate phosphorylation are still stimulated by the substrate, the αG helix likely does not play a significant role in the activation of GCN2 by eIF2α.

While mutation of Thr943 abrogates eIF2α phosphorylation, it does not impair substrate binding to GCN2 (**Fig S6E**), raising the possibility of a second eIF2α binding site within GCN2. Each of the four ISR kinases harbors a loop insertion between the β4 and β5 strands in the N-terminal lobe of the kinase domain (**Fig S7A)**. These loops are predicted to be largely unstructured and show little homology to one another (**Fig S7A, B**). The loop in PERK is rich in S/T residues (**Fig S7B, C**) and when phosphorylated, these residues provide an additional interface for eIF2α binding separate from the canonical αG helix (*40*). This interaction is proposed to improve the efficiency of catalysis by locally concentrating the substrate (*40*).

The human GCN2 kinase domain insertion loop has a lower density of S/T residues but is modestly enriched in D/E residues (**Fig S7B**). These negative charges could recruit eIF2α analogously to the phosphorylated Ser/Thr residues of PERK. If substrate recruitment to this loop stimulates GCN2 activity, autophosphorylation of a variant lacking the loop should be refractory to eIF2α stimulation. Indeed, autophosphorylation of a GCN2 variant lacking a 125 amino acid stretch containing much of the loop (positions 663-788; ΔKDloop) is not stimulated by eIF2α and exhibits reduced inhibition by tRNA (**Fig 5A; Fig S8A, B**), consistent with the insertion loop mediating substrate activation of GCN2. Strikingly, this deletion renders the kinase constitutively active, exhibiting both auto- and substrate phosphorylation even in the absence of P-stalk or ribosome agonists (**Fig 5A**).

**Figure 5.**
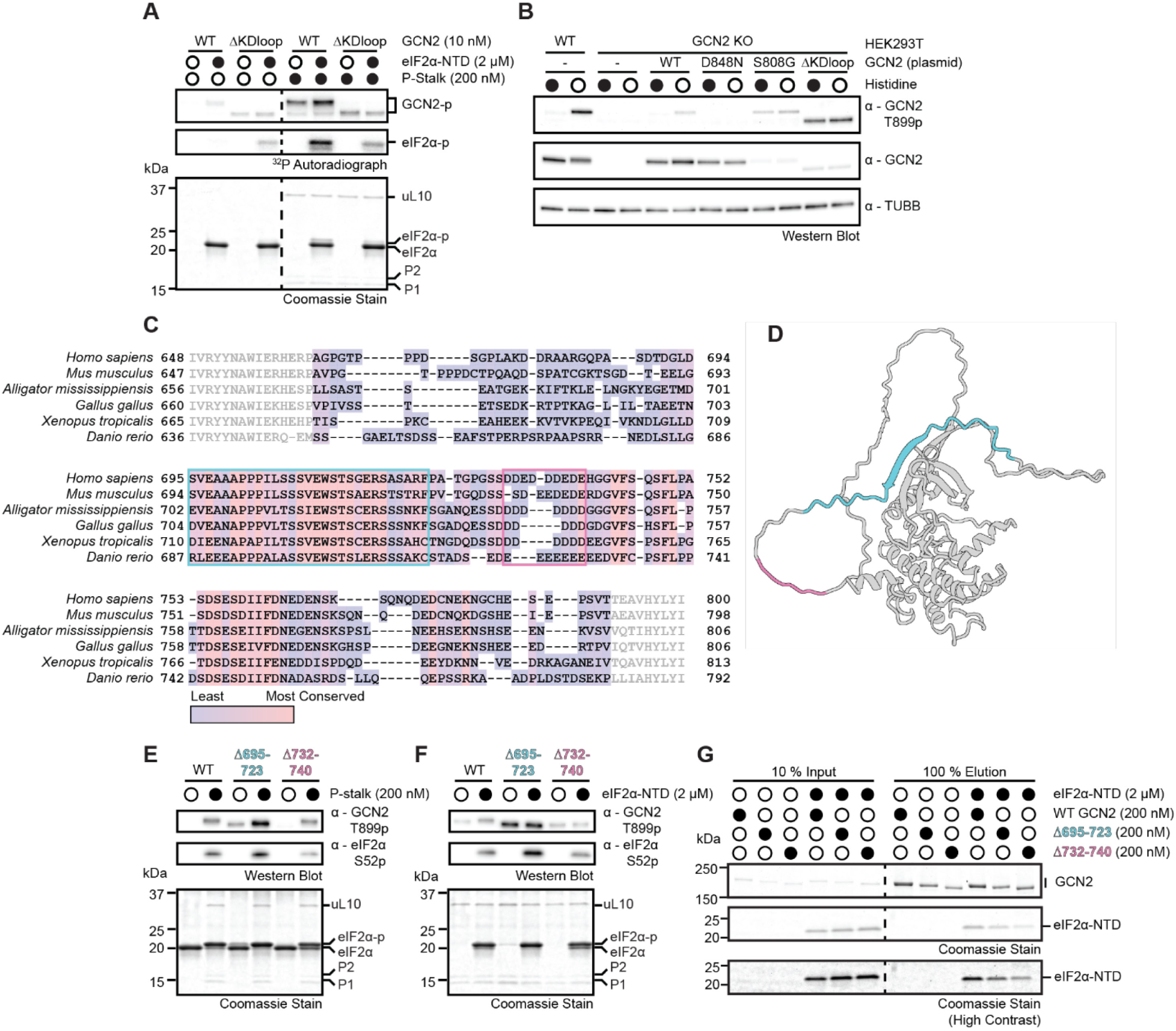
GCN2 Determinants of Activation by Substrate. **(A)** Coomassie stain and autoradiographs of kinase assays with wild-type (WT) and kinase insertion loop deletion (ΔKDloop) GCN2 variants in the presence of 200 nM P-stalk and 2 µM eIF2α-NTD, as indicated. **(B)** Immunoblots of GCN2 activity in wild-type (WT) or ΔGCN2 HEK293T cells transfected with no DNA, wild-type (WT) GCN2 (pMC256), kinase dead GCN2 D848N (pMC574), constitutive active GCN2 S808G (pMC576), or GCN2 ΔKDloop (pMC581) grown on complete media or starved for Histidine for 1 h. **(C)** GCN2 kinase domain insertion loop from species of 6 representative phyla drawn from a Clustal Omega alignment of 818 *Chordata* species. Conservation across all 818 species is indicated in purple to pink color gradient. Insertion loop residues are indicated by darker font. The conserved region containing a predicted β-strand (695-723 in hsGCN2) is boxed in blue. The region containing an acidic patch (732-740) is boxed in pink. **(D)** Structural model of the GCN2 kinase domain (585-1016, *hs*GCN2; Alphafold3). The conserved β-strand and acidic patch are colored as in (C). **(E)** Coomassie stain and immunoblots with the indicated antibodies of kinase assays with 20 nM of the indicated GCN2 variants and 2 µM eIF2α-NTD in the presence or absence of 200 nM P-stalk. **(F)** As in E in the presence or absence of 2 µM eIF2α-NTD. **(G)** Coomassie stains of co-immunoprecipitation experiments with 200 nM of the indicated GCN2 variants in the presence or absence of 2 µM eIF2α-NTD. The region including eIF2α-NTD is displayed at two separate contrast levels for clarity.

This *in vitro* constitutive activity of the KDloop deletion is also observed *in vivo*. Endogenous GCN2 is autophosphorylated at Thr899 upon starvation for amino acids such as Histidine (**Fig 3B**, **5B**). Wild-type GCN2 expressed from a transfected plasmid in a GCN2 KO HEK293T cell line at near endogenous levels also undergoes autophosphorylation upon His starvation, but a transfected kinase-dead D848N variant does not, and the constitutive S808G variant is phosphorylated at Thr899 even in complete media (**Fig 5B**). Remarkably, the ΔKDloop variant also displays activation loop phosphorylation in both starved and unstressed cells (**Fig 5B**), consistent with its high basal activity *in vitro*. Notably, both constitutively active variants are expressed at lower levels than the wild-type or kinase-dead proteins (**Fig 5B**); their basal activity likely induces the integrated stress response which represses their own translation.

Together, these data suggest the GCN2 insertion loop is an auto-repressive element whose effects are relieved by substrate binding. Two motifs within the loop stand out as potential regulatory elements. The first is a highly conserved segment including a β-strand predicted by AlphaFold to complement β4 and extend the 5-stranded β-sheet of the kinase domain N-terminal lobe (positions 695-723; **Fig 5C, D**). The second motif constitutes an acidic patch of consecutive Asp and Glu residues (position 732-740), which is present in all *Chordata* species and comprises the largest negatively charged patch within the loop (**Fig 5C, D**). The GCN2 variant lacking the first motif (Δ695-723) has high constitutive activity and its autophosphorylation is less responsive to both eIF2α and tRNA (**Fig 5E, F; Fig S8C**), suggesting that the conserved β-strand is repressive. In contrast, deletion of the acidic patch (Δ732-740A) renders the kinase less active, implying it is not a negative regulator like the first motif. However, loss of the acidic patch still abrogates both stimulation by the substrate and inhibition of autophosphorylation by tRNA (**Fig 5E, F; Fig S8C**), consistent with a role for this patch in relieving the repression of the neighboring β-strand. Finally, co-immunoprecipitation experiments demonstrate that the acidic patch is required for high affinity binding of eIF2α to GCN2, whereas the conserved β-strand is dispensable (**Fig 5G**). Altogether, these mutagenesis experiments establish the kinase domain insertion loop as a regulatory element that binds eIF2α substrate through its acidic patch to alleviate the repression of the adjacent conserved β-strand motif.

### eIF2α Determinants of GCN2 Activation

Kinases are often regulated by feedback mechanisms whereby phosphorylation of a substrate potentiates the activity of the kinase (*52*). This feedback accelerates the kinases’ response to triggering agonists. The activation of GCN2 by eIF2α could be an example of a direct feedback mechanism to increase potency of the GCN2 response to low levels of ribosome collisions. In this model, a small population of active GCN2 would phosphorylate eIF2α. Phospho-eIF2α could then activate further GCN2 proteins, rapidly amplifying the initial agonism by the ribosome.

GCN2 phosphorylates eIF2α on S52 within an exposed loop spanning positions 47-66 of the S1 domain (**Fig 6A, B**). Were substrate phosphorylation required to stimulate the kinase’s autophosphorylation, then a phospho-null S52A variant of eIF2α would be ineffective, whereas a phosphomimetic S52D variant would more potently activate GCN2. However, the S52A variant instead stimulates GCN2 autophosphorylation equivalently to the WT protein, whereas the S52D variant does not (**Fig 6B, C**), suggesting that direct feedback through substrate phosphorylation does not regulate the activation of GCN2. In fact, eIF2α S52D also binds to GCN2 with lower affinity than the WT or S52A variant (**Fig 6D**), consistent with phosphorylation mediating the substrate’s release from the kinase, not its recruitment.

**Figure 6.**
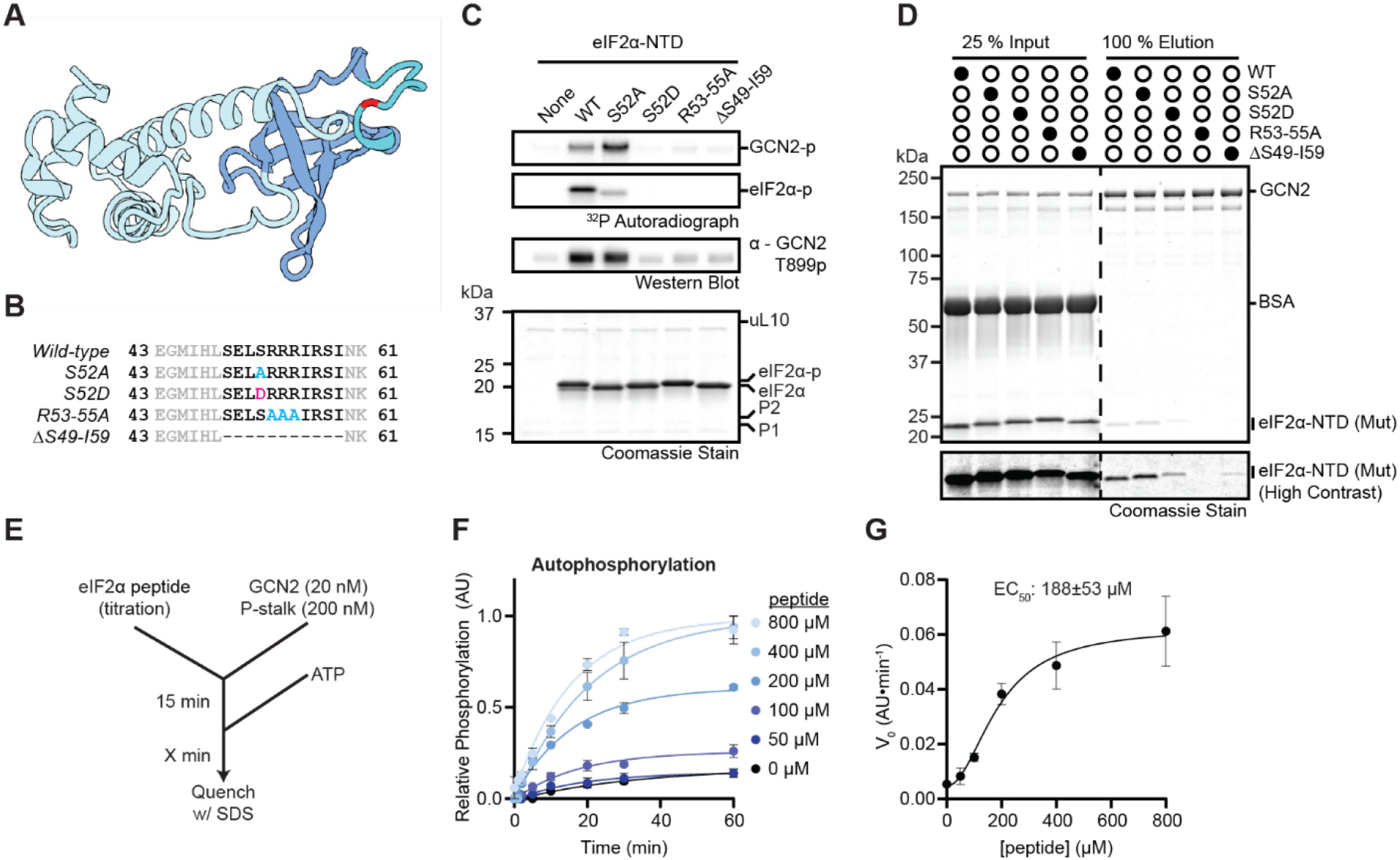
eIF2α Determinants of GCN2 Activation. **(A)** Structural model of the eIF2α-NTD (2-185, *hs*EIF2S1; Alphafold3). The S1 domain is shaded in darker blue, the phosphorylated loop (S49-I59) in cyan, and S52 in red. **(B)** Variants of eIF2α-NTD with Aspartate substitutions colored in magenta and Alanine substitutions colored in cyan. **(C)** Coomassie stain, autoradiographs, and immunoblots with the indicated antibodies of kinase assays with 20 nM GCN2, 200 nM P-stalk, and 4 µM of the indicated eIF2α-NTD variants. **(D)** Coomassie stains of co-immunoprecipitation experiments with 200 nM of GCN2 and 2 µM of the indicated eIF2α-NTD variants. The region including eIFα-NTD is displayed at two separate contrast levels for clarity. **(E)** Flow chart of a kinase experiment as in Fig 4G, but with eIF2α peptide in place of eIF2α-NTD. **(F)** Intensities of autophosphorylation autoradiographs from experiments as in (E) with 200 nM P-stalk (Figure S9C) plotted and fit with single exponential functions. Error bars represent S.E.M. of 2 independent replicates. **(G)** V_0_ in (F) as a function of eIF2α-NTD concentration fit with a 4-term agonist dose response curve (see Methods). Error bars represent S.E.M. of 2 different replicates. Error on reported EC_50_ is the standard error of the fit.

The residues directly following S52 in the eIF2α phosphorylated loop are highly basic (**Fig 6B**). As an acidic patch on GCN2 is required for substrate stimulation, and the introduction of a single acidic residue adjacent to this motif (S52D) is sufficient to disrupt substrate binding and stimulation of GCN2, then neutralization of this basic patch might be predicted to have similar effects. Indeed, mutation of eIF2α R53-55 to Ala oblates both GCN2 binding and stimulation of autophosphorylation (**Fig 6B, C, D**), as does deletion of the entire region from residues 49-59 (**Fig 6B, C, D**). These data suggest that interactions between the basic stretch surrounding S52 and the acidic patch in the GCN2 insertion loop could be responsible for the stimulation of kinase autophosphorylation by eIF2.

In fact, this region alone is sufficient to stimulate GCN2 autophosphorylation. Previous reports have shown that a peptide consisting of the phosphorylated loop, lacking the rest of the S1 domain, can serve as a substrate *in vitro*, albeit with a much higher K_m_ for GCN2 than eIF2α-NTD or eIF2 trimer (*53*). Indeed, GCN2 incubated with P-stalk phosphorylates a synthesized peptide consisting of residues 47-66 of eIF2α (**Fig 6E; Fig S9A, B, C**). This peptide also potentiates the autophosphorylation of GCN2 on its activation loop, albeit with an EC_50_ much higher than for the eIF2α-NTD (**Fig 6F, G; Fig S9C, D, E**). Like with the full NTD, GCN2 stimulation by the substrate peptide requires basic charges adjacent to S52; both a S52p phosphorylated peptide and a charge neutralized R53-55A variant fail to enhance GCN2 autophosphorylation, whereas the S52A variant is partially activating (**Fig S9A, B**). Thus, the unphosphorylated S52 loop of eIF2α is both necessary and sufficient to stimulate the autophosphorylation of GCN2.

## Discussion

Altogether, these data suggest a model whereby the kinase insertion loop of GCN2 functions as an auto-repressive element, likely through the binding of its conserved predicted β-strand to the β-sheet in the N-terminal lobe of the kinase domain. eIF2 binding to this loop, potentially templated by charge interactions between positive residues surrounding S52 of eIF2α and a negatively charged stretch within the insertion loop, releases the autoinhibition, licensing GCN2 activation. RNAs disrupt the binding of eIF2α to this acidic motif and prevent GCN2 autophosphorylation. This regulation is orthogonal to activation by the ribosome or ribosomal P-stalk, which is communicated primarily through the HisRS and CTD domains (**Fig 7**).

**Figure 7.**
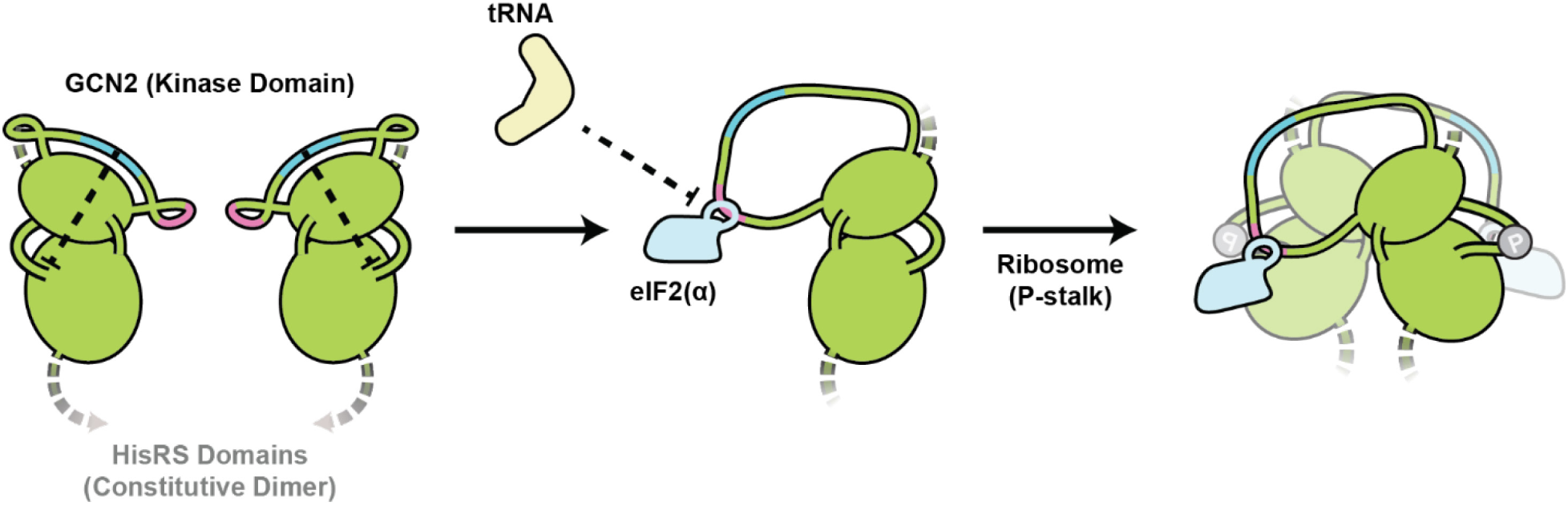
Model of GCN2 Activation by Substrate. GCN2 is constitutively dimerized through interactions in its HisRS and CTD domains. In the absence of substrate and ribosomes, complementation between the conserved β-strand (blue) in the kinase domain insertion loop and β4 in the N-terminal lobe represses GCN2 autophosphorylation. eIF2 binding to the acidic patch displaces the β-strand but is antagonized by tRNA. Once freed from auto-repression, the kinase domains can dimerize in the presence of P-stalk or ribosome to promote autophosphorylation.

The mechanism by which the conserved β-strand within the GCN2 insertion loop represses kinase activity is unclear. Crystallized constructs of the GCN2 kinase domain all lack the flexible insertion loop (*14*, *15*), so its effect on the domain’s conformation is uncertain. A recent study, however, proposed that a GCN2 dimer phosphorylates its activation loop only after parallel back-to-back alignment of the kinase domains mediated in part by a “hydrophobic cradle” formed between the β4s of both N-terminal lobes (*54*). As this is the same interface that the insertion loop β-strand is predicted to bind, an attractive model would be that the insertion loop prevents kinase domain dimerization, and thereby autophosphorylation, until substrate binding to the loop displaces the β-strand and allows interaction between the N-terminal kinase domain lobes (**Fig 7**).

Such allosteric regulation of a kinase by its own substrate is rare, but not unprecedented. For instance, Aurora Kinases B and C (AURKB/C) complex with the centromeric protein INCENP, which is itself an Aurora kinase substrate (*55*). Phosphorylation of the C-terminal “IN box” of INCENP increases the autophosphorylation activity of AURKB/C as well as their activity towards other substrates (*56*, *57*). Likewise, binding of several different G-protein Receptor Kinases (GRKs) to activated GPCRs induces a conformational change to allosterically stabilize the active state of the kinase (*58*, *59*). However, these mechanisms seem distinct from the substrate relief of the auto-repressive β-complementation within GCN2. Instead, the GCN2 β-sheet extension structurally resembles that of the Nuclear Factor κB-Inducing Kinase (NIK), whose N-terminal kinase domain extension contributes two additional β-strands to the kinase domain β-sheet beyond β4(*60*, *61*). However, while the conserved β-strand in GCN2 is inhibitory, the N-terminal extension of NIK is activating, as it packs against the N-terminal kinase lobe and is thought to stabilize its active conformation (*60*, *61*). As such, it seems possible that the mechanism of substrate-mediated activation of GCN2 through relief of its autorepression may be unique to this kinase.

In principle, such substrate-dependent activation could increase the specificity of the kinase for its substrate. A potential GCN2 substrate would have to bind first the insertion loop to activate the kinase and then the active site to be phosphorylated. However, phosphorylation by all four ISR kinases is already considered highly specific for eIF2α, requiring the S1 domain of eIF2α to bind the αG helix of the kinase domain to enforce stringent substrate selection^33,44^. It is unclear why GCN2 would involve an extra layer of regulation, or whether the auto-repression of GCN2 may instead have other roles.

For instance, the GCN2 insertion loop may function as a platform for external regulatory factors to modulate kinase activity. Phosphorylation of the analogous loop in PERK recruits eIF2 to the kinase to increase the local concentration of substrate (*40*). However, competition between other cellular factors and the substrate for these binding sites can regulate and restrict the kinase (*62*). For example, the mitochondrial OMM protein ATAD3A binds this loop and represses PERK in the vicinity of mitochondrial membranes (*63*). This presents a tempting parallel for regulation of GCN2 by RNAs. Though deacylated tRNAs have long been considered agonists of the kinase, they seem largely inert *in vivo* and are repressive *in vitro*. Indeed, if deacylated tRNAs were sufficient to activate GCN2 *in vivo*, they would render the kinase nearly constitutively active as, despite the efficiency of acylation, the enormous abundance of tRNA is predicted to leave around 5 µM tRNA deacylated at any given time, even in cells replete with amino acids (*64*, *65*). Rather, RNAs, including deacylated tRNAs, inhibit GCN2 by blocking eIF2 binding to the GCN2 insertion loop. In cells, this may dampen the basal activity of GCN2, preventing spurious eIF2 phosphorylation.

Endogenously, this repression could be lifted in part by the activities of GCN1, ABCF3 (GCN20 in yeast), or conformational rearrangements of GCN2 induced by binding collided ribosomes. Indeed, the regulation outlined here is only a piece of the emerging picture of GCN2 activation in cells. GCN2 is basally inhibited not only by its own auto-repression and RNAs, but also potentially by translation elongation factors, such as eEF2, which impede its access to the P-stalk, and interactions with free 60S subunits (*26*, *66*). *In vitro*, the absence of these inhibitors permits GCN2 activation independent of GCN1 or ribosome collisions (*29*). However, in cells these positive regulatory factors are likely required to strip away the layers of inhibition and recruit GCN2 to its ideal agonist, a ribosome with an empty A site and an accessible P-stalk (*26*, *30*, *67*).

It is also possible that RNAs exert their inhibitory activity *in vitro* not by binding to GCN2, but by binding to eIF2α. The GTP-bound eIF2 trimer binds tRNA_iMet_ with high affinity and specificity. However, in the absence of GTP and tRNA_iMet_, the complex can bind other RNAs with lower affinity (*68*), as can the isolated α subunit (*69–71*). If promiscuous RNA binding to eIF2α displaced it from the GCN2 insertion loop *in vitro*, this would manifest as inhibition of GCN2 by RNA. In cells, the eIF2 trimer exists in both a ternary complex bound with tRNA_iMet_ and GTP, as well as a tRNA-free GDP-bound form. If tRNA-bound eIF2 is inhibitory *in vitro*, this implies that the tRNA_iMet_-bound ternary complex may be an insufficient co-activator of GCN2 *in vivo* and that the GDP-bound eIF2 trimer lacking tRNA would instead be a preferred substrate. As phosphorylated eIF2 inhibits eIF2B, its own GEF required for tRNA binding, this would produce a feedback mechanism whereby continued activity of GCN2 and inactivation of eIF2B builds up an increasingly larger pool of tRNA-free eIF2, which in turn further activates GCN2.

Clearly, the role of the substrate as an active participant in GCN2 activation has been underappreciated. While further studies will establish the ramifications for this regulation *in vivo*, the careful *in vitro* reconstitution here has demonstrated that GCN2 is basally autoinhibited by its kinase domain insertion loop and that allosteric activation by its own substrate relieves this repression to unleash full kinase activity. The kinase insertion loop also offers an alternative pharmacological target to inhibit or agonize the kinase *in vivo* (*62*). Existing ISR inhibitors tend to suffer from “paradoxical activation” whereby high concentrations of these ATP-competitive drugs activate related kinases (*72*, *73*). Peptides or small molecules that disrupt or reinforce the binding between eIF2 and GCN2 could inactivate or potentiate GCN2, respectively, without perturbing the activity of the other ISR kinases, which appear to lack an auto-repressive insertion loop. Considering the disease implications of ISR dysregulation, such meticulous control of a kinase poised to impact cell fate seems appealing. Its discovery here represents an important step in understanding control of GCN2 and the ISR and introduces a new paradigm for kinase regulation in general.

## Acknowledgements

We thank Dr. Chien-I Yang for the PKR KO cell line; Dr. Roger Williams for the P-stalk expression plasmid; Dr. Frances Diehl for the GFP expression plasmid; and Dr. Eugene Park and Dr. Frances Diehl for their critical reading of the manuscript.

## Author Contributions

M.A.C: Conceptualization, Methodology, Investigation, Formal Analysis, Writing – original draft, Writing – review and editing; R.G. Conceptualization, Resources, Supervision, Writing – review and editing, Funding Acquisition.

## Data, code, and materials availability

All materials are available upon request from RG.

## Supplementary Materials

## Materials and Methods

### DNA Constructs

#### Overexpression Plasmids for Protein Purification

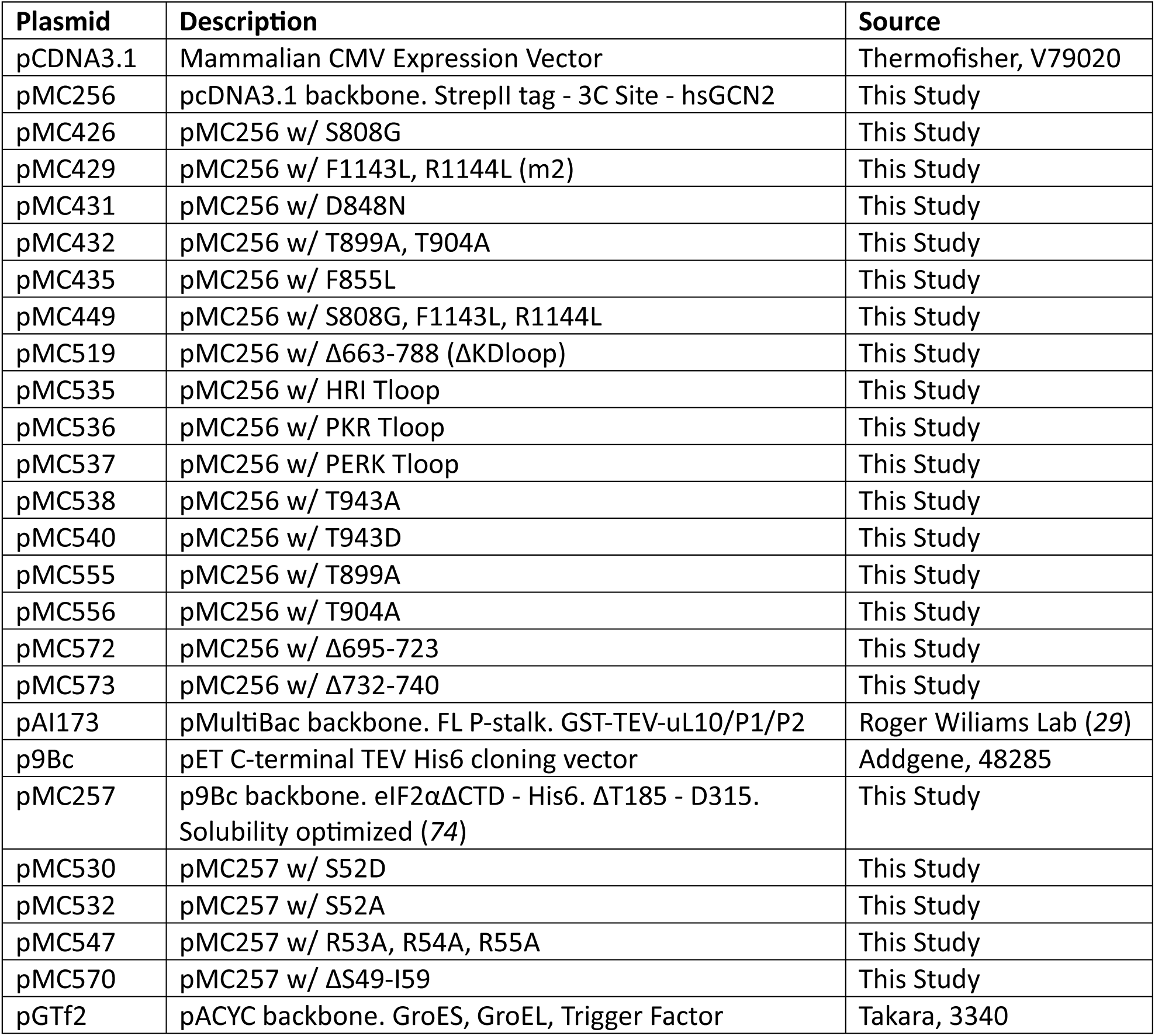

#### Transfection Plasmids for In vivo Experiments

| Plasmid | Description | Source |
| --- | --- | --- |
| pIRE4-Azi | Synthetase tRNA pair for Azido-phenylalanine (AzF) incorporation at amber codons in mammals | Addgene, 105829 |
| pANAP | Synthetase tRNA pair for Anap incorporation at amber codons in mammals, | Addgene, 48696 |
| pMC389 | pIRE4-Azi w/ Synthetase ORF deleted | This Study |
| pMC390 | pANAP w/ Synthetase ORF deleted | This Study |
| pFN24K | pFN24K Mammalian strongly compromised CMV Expression Vector | Promega, G3780 |
| pMC505 | pFN24K backbone. 3xFlag tag - hsGCN2 | This Study |
| pMC574 | pMC505 w/ D848N | This Study |
| pMC576 | pMC505 w/ S808G | This Study |
| pMC581 | pMC505 w/ Δ663-788 | This Study |
| pFD042 | pFC15K backbone. eGFP - 3C - StrepII - 3xFlag Tag | Frances Diehl (75) |

#### CRISPR Knockout Plasmids

| Plasmid | Description | Source |
| --- | --- | --- |
| pX330 | pX330-U6-Chimeric_BB-CBh-hSpCas9 | Addgene, 158973 |
| pMC262 | pX330 backbone. GCN2 sgRNA1, Ex9 nt1120 | This Study |

#### In vitro Transcription Oligos

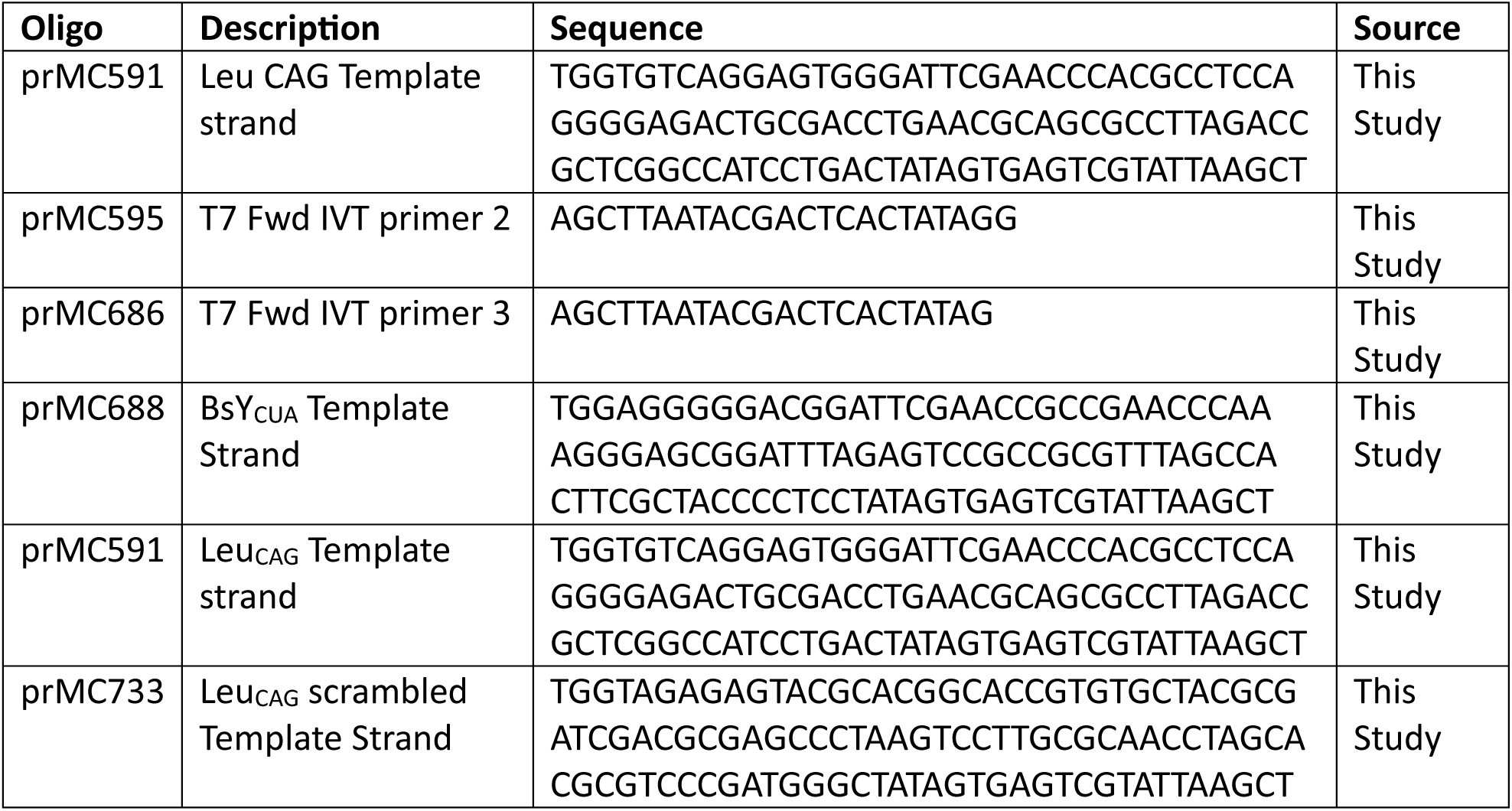

#### Northern Probes

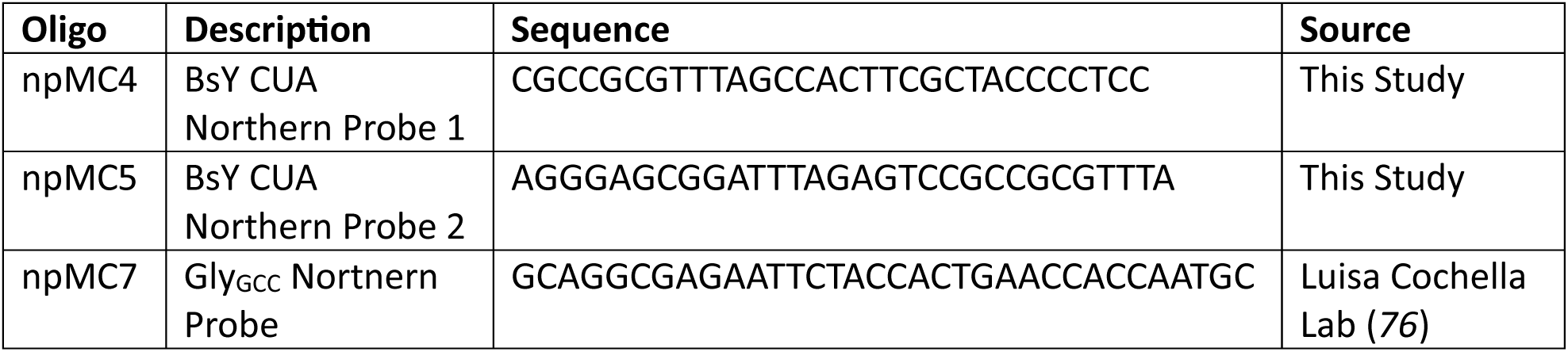

### Antibodies

#### Primary Antibodies

**Antibodies** *Primary Antibodies*
| Antibody | Dilution | Company | Catalog # |
| --- | --- | --- | --- |
| Rabbit Monoclonal $\alpha$ - Total GCN2 | 1:2000 | Cell Signaling Technology | 46041 |
| Rabbit Monoclonal $\alpha$ - GCN2 (T899p) | 1:1000 | Cell Signaling Technology | 94668 |
| Mouse Monoclonal $\alpha$ - Total eIF2α | 1:2000 | Abcam | ab5369 |
| Rabbit Monoclonal $\alpha$ - eIF2α (S52p) | 1:1000 | Abcam | ab32157 |
| Rabbit Polyclonal $\alpha$ - uS3 | 1:5000 | Abcam | ab140688 |
| Rabbit Polyclonal $\alpha$ - eRF1 | 1:1000 | Cell Signaling Technology | 13916 |
| Rabbit Monoclonal $\alpha$ - Tubulin $\beta$ | 1:2000 | Cell Signaling Technology | 2128 |
| Rabbit Polyclonal $\alpha$ - GFP | 1:5000 | Abcam | ab6556 |
| Mouse Monoclonal $\alpha$ - Vinculin | 1:5000 | Santa Cruz Biotechnology | sc-73614 |

#### Secondary Antibodies

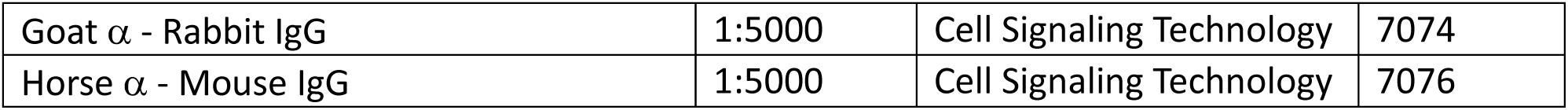

### Protein Purification

#### GCN2

Human GCN2 was overexpressed in Expi293F (Thermofisher, A14527) by transfection of plasmid encoding wild-type hsGCN2 or mutants (see DNA Constructs). Cells were cultured in Expi293 Expression Medium (Thermofisher, A1435101). The day before transfection cells were subcultured to a density of 2.5-3E6 cells/ml in 25 ml. On the day of transfection cells were diluted with fresh media to a density of 2.5E6 in 25 ml and transfected with 25 µg plasmid using Expifectamine293 (Thermofisher, A14524) according to the manufacturers protocol. 16 hours after transfection, ExpiFectamine™ 293 Transfection Enhancers 1 and 2 (Thermofisher, A14524) were added to the media, along with 2 µM GCN2 inhibitor A-92 (MedChemExpress, 1448693-69-3) and 500 nM ISRIB (Millipore Sigma, SML0843-5MG) to suppress activity of constitutively active mutants. Protein expression continued for 48 hours before cells were harvested by centrifugation at 300 x *g* for 5 min at 25°C and flash frozen in liquid nitrogen.

To purify the expressed protein, pellets were thawed on ice and resuspended in 25 ml of Buffer A (20 mM Tris·HCl pH 8.0, 150 mM NaCl, 5 % Glycerol, 2 mM β-mercaptoethanol (BME)). Resuspended pellets were supplemented with 1 tablet cOmplete EDTA-free protease inhibitor cocktail (Millipore Sigma, 4693159001), 500 U Benzonase Nuclease (Millipore Sigma, E8263-5KU), and 800 µl BioLock reagent (IBA LifeSciences, 2-0205-050). Cells were lysed by sonication using a probe sonicator with a ½” diameter horn (Branson Ultrasonics, SFX250) using a 1 min cycle at 50 % amplitude with 3 sec pulses followed by 9 second rests. The resulting lysate was clarified by centrifugation at 30,000 x *g* for 30 min at 4°C.

The clarified lysate was syringe filtered with a 0.45 µm SFCA filter (Corning, 431220) and loaded at 1 ml/min onto a 5 ml Strep-Tactin 4Flow high capacity FPLC column (IBA LifeSciences, 2-1258-001) equilibrated with 4 column volumes (C.V.) of Buffer A. The bound protein was washed with 10 C.V. of Buffer A before elution with 5 C.V. of Buffer A supplemented with 6 mM Desthiobiotin (IBA LifeSciences, 2-1000-002). The eluted protein was immediately loaded at 1 ml/mi onto a 5 ml HiTrap Q HP anion exchange column (Cytiva, 17115301) in equilibrated in Buffer A. The bound protein was washed with 5 C.V. of Buffer A at 2 ml/min flow and eluted with a 2 C.V. gradient from 100 % Buffer A to 100 % Buffer B (20 mM Tris·HCl pH 8.0, 1000 mM NaCl, 5 % Glycerol, 2 mM BME) under low flow, at 0.25 ml/min. The elution peak containing GCN2 was generally less than 1 ml and could be loaded directly onto a Superdex Increase S200 10/300 GL size exclusion chromatography column (Cytiva, 28990944) equilibrated in Buffer C (20 mM HEPES pH 7.5, 150 mM NaCl, 5 % Glycerol, 2 mM Tris(2-carboxyethyl)phosphine hydrochloride (TCEP)). Peak fractions were collected, diluted to 1 µM in Buffer C, and flash frozen in liquid nitrogen.

#### P-stalk

P-stalk heteropentamer was expressed in insect cells and purified based on published protocols (*29*). P-stalk bacmid encoding each of GST-TEV-uL10, RPLP1, and RPLP2 were generated by recombination between a donor plasmid (DNA constructs) and bacmid maintained in MAX Efficiency DH10Bac Competent Cells (Thermnofisher, 10361012) using the Bac-to-bac system (Thermofisher, 10359016) according to the manufacturer’s protocols. Recombinant bacmid DNA was purified using PureLink HiPure miniprep kits (Thermofisher, K210002) and transfected into Expisf9 Cells (Thermofisher, A35243) grown to 2.5E6 cells/ml in 4 ml ExpiSf CD Medium (Thermofisher, A3767802) using Expifectamine Sf (Thermofisher, A38915) according to the manufacturer’s protocol. Transfected cells were grown for 4-5 days until their viability dropped below 80 %. The resulting P0 baculovirus was harvested from the media by centrifugation at 500 x *g* for 5 min at 25 °C and stored at 4 °C. Virus was amplified by reinfecting 50 ml of cells grown to 2E6 cells/ml with 10 ul of P0 virus. Cell growth was monitored from 4-7 days and P1 virus was harvested as before when cell viability dropped below 80 %.

P-stalk proteins were expressed by infecting 50 ml of ExpiSf9 cells grown to 5E6 cells/ml with 1 ml of P1 24 hours after the addition of 1 ml of ExpiSf Enhancer (Thermofisher, A3767806). Expression continued for 72 hours before cells were harvested by centrifugation at 300 x *g* for 5 min and flash frozen in liquid nitrogen.

To purify the expressed protein, pellets were thawed on ice and resuspended in 50 ml of Buffer D (20 mM Tris·HCl pH 8.0, 100 mM NaCl, 5 % Glycerol, 5 mM Dithiothreitol (DTT)). Resuspended pellets were supplemented with 1 tablet cOmplete EDTA-free protease inhibitor cocktail (Millipore Sigma, 4693159001) and 500 U Benzonase Nuclease (Millipore Sigma, E8263-5KU). Cells were lysed by sonication using a probe sonicator with a ½” diameter horn using a 1 min cycle at 50 % amplitude with 3 sec pulses followed by 9 second rests. The resulting lysate was clarified by centrifugation at 30,000 x *g* for 30 min at 4°C.

The clarified lysate was syringe filtered and loaded onto a 5 ml GSTrap FF FPLC column (Cytiva, 17513101) equilibrated in 4 C.V. Buffer D. The bound protein was washed with 10 C.V. of Buffer D and incubated overnight with 5 ml of 30 µg/ml TEV protease diluted in 5 ml Buffer D on the column. The next morning the cleaved protein was collected from the column and loaded at 1 ml/mi onto a 5 ml HiTrap Q HP anion exchange column in equilibriated in Buffer D. The bound protein was washed with 5 C.V. of Buffer D at 2 ml/min flow and eluted with a 10 C.V. gradient from 100 % Buffer D to 100 % Buffer E (20 mM Tris·HCl pH 8.0, 1000 mM NaCl, 5 % Glycerol, 5 mM DTT) at 1 ml/min. Peak fractions containing P-stalk proteins were concentrated to < 500 µl and loaded onto a Superdex Increase S200 10/300 GL size exclusion chromatography column equilibrated in Buffer F (20 mM HEPES, 50 mM KOAc, 5 mM Mg(OAc)_2_, 5 % Glycerol, 2 mM TCEP). Peak fractions containing P-stalk proteins were collected, concentrated to 20 µM and flash frozen in liquid nitrogen.

#### eIF2α-NTD

eIF2α-NTD was expressed in bacterial cells and purified using on published protocols (*74*). BL21 *E. coli* (New England Biolabs, C2527H) were co-transformed with plasmids encoding eIF2α-NTD-His6 (DNA constructs) and pGTf2 (Takara, 3340), to express chaperones GroEL and Trigger Factor, and grown overnight at 37 °C in LB media supplemented with 100 ug/ml Carbenicillin and 25 ug/ml Chloramphenicol. Overnight cultures were diluted 1:100 into 1L of LB medium supplemented with the same antibiotics and grown at 37 °C. Chaperone expression was induced at OD 0.4 by addition of 10 ng/ml tetracycline. At OD 0.6, cultures were chilled to 20 °C, and eIF2α-NTD expression was induced by addition of 0.5 mM Isopropyl β-D-1-thiogalactopyranoside (IPTG, Gold Biotechnology, I2481C25). Expression continued for 6 hours at 20 °C before cells were harvested by centrifugation for 10 min at 4000 x *g* and flash frozen in liquid nitrogen.

The cell pellet from 2 L of culture was thawed on ice, resuspended in 50 ml of Buffer G (20 mM Tris·HCl pH 8.0, 300 mM NaCl, 25 mM Imidazole, 5 % Glycerol, 2 mM BME) supplemented with 1 tablet cOmplete EDTA-free protease inhibitor cocktail and 500 U Benzonase Nuclease, and lysed by sonication using a probe sonicator with a ½” diameter horn using a 2 minute cycle at 60 % amplitude with 3 sec pulses followed by 9 second rests. The resulting lysate was clarified by ultracentrifugation using a Ti45 rotor (Beckman Coulter, 339160) for at 40,000 rpm for 45 min at 4 °C.

The clarified supernatant was added to 1 ml of PureCube Ni-NTA agarose resin (Cube Biotech, 31103) equilibrated w/ 10 C.V. of Buffer G. The lysate was mixed with the resin for 1 h with gentle rotation at 4 °C and poured into a chromatography column. The unbound flow-through was removed and the beads were washed with 10 C.V, of Buffer G, followed by 10 C.V. of Buffer H (20 mM Tris·HCl pH 8.0, 1000 mM NaCl, 25 mM Imidazole, 5 % Glycerol, 2 mM BME) and a final 10 C.V. of Buffer G. Proteins bound after washing were eluted by 5 1 C.V. additions of Buffer I (20 mM Tris·HCl pH 8.0, 300 mM NaCl, 250 mM Imidazole, 5 % Glycerol, 2 mM BME), concentrated to < 1 ml, and loaded onto a Superdex 75 10/300 GL size exclusion column (Cytiva, 29148721) equilibrated in Buffer J (20 mM HEPES pH 7.5, 350 mM NaCl, 5 % Glycerol, 2 mM TCEP). Peak fractions were collected, concentrated to 100 µM, and flash frozen in liquid nitrogen.

#### Endogenous eIF2

Endogenous eIF2 trimer was purified from Expi293F cells based on published protocols (*77*, *78*). 400 ml of Expi293 cells were grown to a density of 5E6 cells/ml in Expi293 Expression Medium. The cells were pelleted by centrifugation at 2,500 x *g* for 15 min at 4 °C, washed in 40 ml ice cold 1X PBS (Thermofisher, 10010023), and pelleted again by centrifugation at 2,500 x *g* for 5 min at 4 °C. The washed pellet was resuspended in 25 ml Buffer K (10 mM HEPES pH 7.5, 10 mM KCl, 1.5 mM MgCl_2_, 1 mM DTT, 1 X Halt Protease/Phosphatase Inhibitor Cocktail (Thermofisher, 78440)) and immediately pelleted by centrifugation at 2,500 x *g* for 5 min at 4°C. The pellet was resuspended in 15 ml ice cold Buffer K and incubated on ice for 10 min before being lysed by 10 strokes with the tight pestle of a 40 ml Kimble dounce homogenizer (Millipore Sigma, D9188-1SET). Unbroken nuclei were removed from the lysate by centrifugation at 3,000 x *g* for 20 min at 4 °C and the resulting supernatant was supplemented with 10 % Glycerol, 1 mM EDTA, and 1mM EGTA before clarification by centrifugation at 20,000 x *g* for 20 min at 4 °C. After clarification, the KCl concentration was adjusted to 450 mM and the lysate was ultracentrifuged at 45,000 rpm for 4 h at 4 °C in a Ti70 rotor (Beckman Coulter, 342184). The middle 2/3 of the supernatant was carefully extracted and stirred in a beaker at 4 °C for 1 h as saturated (NH_4_)_2_SO_4_ was slowly added to a final concentration of 40 %. Precipitated proteins were removed by centrifugation at 20,000 x *g* for 10 min at 4 °C. The supernatant was again added to a beaker and stirred at 4 °C for 1 h as saturated (NH_4_)_2_SO_4_ was slowly added to a final concentration of 50 %. Precipitated proteins were again pelleted by centrifugation at 20,000 x *g* for 10 min at 4 °C, and the pellet containing eIF2 was resuspended in 20 ml Buffer L (20 mM HEPES pH 7.5, 50 mM KCl, 10 % Glycerol, 1 mM DTT) and dialyzed overnight against 2 L of Buffer L.

The next morning, the protein was recovered and syringe filtered with a 0.45 µm SFCA filter before loading onto a monoQ 5/50 anion exchange column (equivalent to Capto HiRes Q 5/50, Cytiva, 29275878) equilibrated in Buffer L. Proteins washed with 5 C.V. of Buffer L and eluted in a 20 ml gradient from 100 % Buffer L to 100 % Buffer M (20 mM HEPES pH 7.5, 500 mM KCl, 10 % Glycerol, 1 mM DTT), and fractions containing eIF2 were pooled and dialyzed for 2 h against 2L of Buffer L. The protein was again syringe filtered and subjected to identical ion exchange chromatography protocol using a MonoS 5/50 cation exchange column (equivalent to Capto HiRes S 5/50, Cytiva, 29275877). Fractions containing eIF2 were pooled and concentrated to < 500 µl before loading onto a S200 increase size exclusion column equilibrated in Buffer N (20 mM HEPES pH 7.5, 150 mM KCl, 10 % Glycerol, 1 mM DTT). Peak fractions containing eIF2 were pooled, concentrated to 2.5 µM, and flash frozen in liquid nitrogen.

### RNA Purifications

#### Ribosomes

80S ribosomes were purified from rabbit reticulocyte lysate (RRL, Green Hectares). 6 ml of crude RRL was split 4 ways, layered 1.5 ml per tube over 1 ml sucrose cushions made of Buffer O (50 mM Tris·HCl pH 8.0, 500 mM KOAc, 25 mM Mg(OAc)_2_, 1 mm DTT, 500 mM Sucrose) in 3.5 ml open-top polycarbonate centrifuge tubes (Beckman Coulter, 349622), and centrifuged at 100,000 x *g* for 1 h at 4 °C in a TLA100.3 rotor (Beckman Coulter, 349481). The resulting pellets were resuspended in 1.5 ml total volume of Buffer P (50 mM HEPES pH 7.5, 500 mM KCl, 2 mM MgCl_2,_ 2 mM DTT). To separate the ribosomal subunits, 2 mM Puromycin Dihydrochloride (Millipore Sigma, P7255-25MG) was added to the resuspension, and the ribosomes were incubated sequentially on ice for 15 min, at 37 °C for 10 min with gentle agitation, and on ice for 10 min. This mixture was then layered, 250 µl per tube, above 12 ml gradients from 10 – 35 % sucrose made in Buffer P with a Biocomp Gradient Master (Biocomp Instruments, Model 108). The gradients were centrifuged at 40,000 rpm for 1:45 at 4 °C in open-top 12 ml PolyClear tubes (Seton Scientific, 7030) using a Sw41 Ti swinging bucket rotor (Beckman Coulter, 331362). The gradients were fractionated into 20 500 µl fractions using a piston fractionator (Biocomp Instruments, Model 152) with continual Absorbance_260_ monitoring using a Triax flow cell (Science Services, FC-2). Fractions containing 40S and 60S subunits were pooled and dialyzed overnight against 2 L Buffer Q (20 mM HEPES pH 7.5, 100 mM KOAc, 5 mM Mg(OAc)_2_, 2 mM DTT).

The next day, the dialyzed ribosomes were concentrated to < 1.5 ml using a 15 ml 100 kDa centrifugal concentrator (Millipore Sigma, UFC910008). The resulting concentrate was layered, 250 µl per tube, above 12 ml gradients from 10 – 35 % sucrose made in Buffer Q with a Biocomp Gradient Master. The gradients were centrifuged at 40,000 rpm for 1:45 at 4 °C in open top 12 ml PolyClear tubes using a Sw41 Ti swinging bucket rotor. The gradients were fractionated into 20 500 µl fractions using a piston fractionator. Fractions containing 80S monosomes were pooled and dialyzed overnight against 2 L of Buffer Q. The next day the ribosomes were concentrated to 2 µM using a 15 ml 100 kDa centrifugal concentrator and flash frozen in liquid nitrogen.

#### Bulk tRNA

Total tRNA was purified from pig liver. 600 g pig liver (Pel-Freez, 59423-2) was cut into pieces and homogenized in 1 L ice-cold Buffer R (20 mM Tris·HCl pH 7.5, 100 mM KCl) using a Waring blender (Waring Laboratory Science, 700S). The homogenate was centrifuged at 9,500 rpm for 16 min at 4 °C using a FiberLite F9-6 x 1000 Rotor (Thermofisher, 096-061075) with 1 L PPCO bottles (Thermofisher, 010-1491) resistant to 50 % phenol. The resulting supernatant was mixed with an equal volume phenol (Millipore Sigma, P9346) and stirred at room temperature for 30 min. The mixture was again centrifuged at 12,000 rpm for 15 min at room temperature to separate the phases. The aqueous phase was collected in a new flask and stirred with equal volume chloroform (Millipore Sigma, C2432-1L) for 30 min at room temperature. The mixture was again centrifuged at 12,000 rpm for 15 min. The resulting aqueous phase was collected, mixed with 0.1 volume of 3M NaOAc pH 5.0 and 3 volumes of 95 % ethanol, and stored at -20 °C overnight.

The next morning, the precipitate was centrifuged at 8, 000 rpm for 20 min at 4°C and the resulting crude RNA pellet was washed with 70 % ethanol, air dried and resuspended in 200 ml Buffer R and centrifuged one final time at 8, 000 rpm for 20 min at 4°C to remove any insoluble contaminants. The crude RNA was then loaded onto a 17 ml column packed DEAE Sepharose fast flow resin (Cytiva, 17070960) equilibrated in Buffer R. The column was washed with > 200 ml of Buffer R until the A_260_ of the flow-through dropped below 0.3. The bound tRNAs were eluted in 200 ml Buffer S (20 mM Tris·HCl pH 7.5, 100 mM KCl, 1 M NaCl), mixed with 400 ml 95 % ethanol, and stored at -20 °C overnight. The following morning the precipitate was centrifuged at 8,000 rpm for 25 min at 4 °C. The pellet was washed with 70 % ethanol, air dried, resuspended in Ambion Nuclease Free water (Thermofisher, AM9932) at 10 mg/ml, and flash frozen in liquid nitrogen.

These total tRNA were considered partially acylated and were chemically deacylated by alkaline hydrolysis. 10 mg of total tRNA were thawed on ice, mixed with 0.1 volume of 3M NaOAc pH 5.0 and 3 volumes of 95 % ethanol, and incubated at -20 °C for 1 h. The resulting precipitate was centrifuged at 20,000 x g for 30 min at 4 °C, washed twice with 70 % ethanol, and air dried. The pellet was resuspended in 950 ul nuclease free water, and the pH raised by addition of 50 µl of 1 M Tris·HCl pH 9.0. The tRNA were incubated at pH 9.0 for 1 h at 37 °C and then purified by a second ethanol precipitation. The final pellet was resuspended at 10 mg/m in nuclease-free water and flash frozen with liquid nitrogen. These tRNAs were considered deacylated throughout this study.

#### In vitro transcribed tRNAs

tRNAs and scrambled RNAs were transcribed with the MegaShortScript T7 transcription kit (Thermofisher, AM1354) according to the manufacturers protocol. Single-stranded template DNA oligos including both the T7 promoter and tDNA sequence (DNA constructs) were annealed with ssDNA oligos encoding the T7 complement sequence at 2.5 µM final concentration in Buffer T (10 mM Tris·HCl pH 8.0, 100 mM NaCl, 1 mM EDTA). The oligos were fully denatured by heating to 95 °C for 3 min and then annealed by gradual cooling to room temperature. 50 µl reactions were assembled from kit components with 250 nM annealed oligos, 7.5 mM each of ATP, CTP, GTP, UTP, 1 X T7 Reaction Buffer, and 5 µl T7 Enzyme Mix. Reactions were incubated at 37 °C overnight. The next morning, the DNA template was degraded by addition of 5 U TURBO DNase (Thermofisher, AM2238) at 37 °C for 15 min. Transcription products were purified using RNA Clean and Concentrate columns (Zymo Research, R1019), eluted in 200 µl nuclease-free water, and flash frozen in liquid nitrogen at 1 mg/ml.

### Cell Culture

HEK293T cells (ATCC, CRL-3216) were cultured in Dulbecco’s Modified Eagle Medium (DMEM; Thermofisher, 11995065) supplemented with 10 % fetal bovine serum (FBS; Thermofisher, 26140079) in 10 cm cell culture dishes (Corning, 430167) at 37 °C in a 5 % CO_2_ humidified incubator and passaged every 2-3 days. HEK293T cells were thawed and passaged at least twice before their use in experiments. GCN2 KO cells were produced from parental HEK293T cells by transfection of pMC262 (DNA constructs). Cells were grown for 48 hours post-transfection before dilution with fresh media to single cells in 96-well plates (USA Scientific, CC7682-7596). Cells were outgrown for 14 days and colonies validated as GCN2 knockouts by western blotting and PCR. Positive colonies were selected for clonal expansion and cryogenically stored. PKR KO cells, a gift from Chien-I Yang, were similarly generated from parental HEK293T cells.

### Transfection

tRNA transfections in Figure 3 were performed in PKR KO cells. GCN2 transfections in Figure 5 were performed in GCN2 KO cells. Cells were seeded at 2E6 cells/plate 24 hours prior to transfection and supplied with fresh media 1 hour before transfection. Cells were transfected with the indicated plasmids (DNA Constructs) using Lipofectamine 3000 (Thermofisher, L3000001) according to the manufacturer’s protocol. 5 µg plasmid DNA was transfected per 10 cm dish. No DNA transfection controls were processed identically, including transection reagents but omitting DNA.

### Cellular Treatment and Lysis

For Histidine starvation experiments, DMEM was removed by gentle aspiration 16 – 24 hours after transfection and the cells were washed once with 1X PBS before introduction of DMEM -Histidine Media (USBiologicals, D9801-02). After 1 h, the medium was removed by gentle aspiration and cells were washed rapidly in 1X PBS and harvested by scraping in 250 µl Buffer U (50 mM HEPES pH 7.4, 100 mM KOAc, 15 mM Mg(OAc)_2_, 5 % Glycerol, 0.5 % Triton X-100 (Millipore Sigma, X100-100ML)) supplemented with 1X Halt Protease/Phosphatase Inhibitor Cocktail, 80 U TURBO DNase. Lysates were clarified by centrifugation at 8,000 x *g* for 5 min at 4°C and either flash frozen in liquid nitrogen or directly used for western blotting or sucrose gradient sedimentation.

SRI-41315 experiments were performed in a similar manner. 16-24 hours after transfection, the cells were supplied with fresh media. After an hour, SRI-41315 (Millipore Sigma, SML3312-5MG) was added to a final concentration of 3.3 or 10 µM, as indicated. Control cells were treated with an equal volume of DMSO (Thermofisher, D12345). Cells were cultured for the indicated time in the presence of the drug before being washed rapidly in 1X PBS and harvested by scraping in 250 µl Buffer U (50m M HEPES pH 7.4, 100mM KOAc, 15mM Mg(OAc)_2_, 5 % Glycerol, 0.5 % Triton X-100 (Millipore Sigma, X100-100ML), 1 X Halt Protease/Phosphatase Inhibitor Cocktail, 80 U TURBO DNase. Lysates were clarified by centrifugation at 8,000 x *g* for 5 min at 4°C and either flash frozen in liquid nitrogen or directly used for western blotting or sucrose gradient sedimentation.

### Kinase Assay

Proteins and RNAs were thawed on ice and diluted to 10 X assay concentrations in Buffer V (20 mM HEPES pH 7.4, 50 mM KOAc, 5 mM Mg(OAc)_2_. Reactions were assembled in Buffer V with 0.5 mM TCEP, and 0.5 mg/ml recombinant albumin (NEB, B9200S), and the indicated proteins, RNAs, polyinosine-polycytidylic acid (polyIC; InvivoGen, tlrl-picw), or polyadenylic acid (polyA; Millipore Sigma, 10108626001). Typical reaction conditions included 20 nM GCN2, 2 µM eIF2α-NTD, 200 nM P-stalk, and 5 µM tRNA, though many variations were performed as outlined in the figures. Single timepoint reactions were assembled to a final volume of 9 µl/reaction. Reactions for kinetic experiments were mixed at the same proportions such that 10 µl aliquots could be removed at each time point, with 10 µl remaining unused at the end. Once assembled, reactions were moved to 17 °C and incubated without ATP for 15 min. After this pre-incubation, the reaction was supplemented 500 µM (final concentration) ATP and 0.25 µCi/µl γ^32^P-labeled ATP (Revvity, BLU502A001MC), bringing the final volume to 10 µl/reaction, and returned to 17 °C. At the indicated time points, 9 µl reaction was removed and quenched with 3 µl 4 X Laemmli Buffer (200 mM Tris·HCl pH 6.8, 8 % Sodium Dodecyl Sulfate, 0.4 % Bromophenol blue, 40 % Glycerol).

### Immunoprecipitations

Proteins and RNAs were thawed on ice. 50 µl reactions were assembled with the indicated proteins and RNAs in Buffer V with 0.5 mM TCEP, 0.5 mg/ml recombinant albumin, and 0.1 % Tween-20 (Millipore Sigma, P1379-25ML). Typical reactions contained 200 nM GCN2, 2 µM eIF2α-NTD, 2 µM P-stalk, 0 – 5 µM tRNA, though other variations were tested, as indicated. Reactions were incubated at 17 °C for 15 min, and then 40 µl was added to 0.2 µl MagStrep Strep-Tactin XT beads (IBA Lifesciences, 2-5090-010) equilibrated in Buffer V supplemented with 0.1 % Tween-20, with the rest reserved as an input sample. Reactions were incubated with beads for 1 hour at room temperature and then the unbound fraction was removed using a magnetic tube rack (Zymo Research, 3DP-1002). Beads were washed 3 times with Buffer V supplemented with 0.5 mM TCEP, 0.05 mg/ml recombinant albumin, and 0.1 % Tween-20 and three times with Buffer W (100 mM Tris·HCl pH 8.0, 200 mM NaCl, 1 mM EDTA, 0.1 % Tween-20) before elution in 20 µl Laemmli Buffer or by sequential 15 min incubations with 10 µl Buffer Y (100 mM Tris·HCl pH 8.0, 150 mM NaCl, 1 mM EDTA, 50 mM biotin (IBA Lifesciences, 2-1016-005)).

### Sucrose Gradient Sedimentation

Sucrose gradients were prepared from stocks of 10 % and 50 % sucrose in Buffer Z (25 mM HEPES pH 7.5, 100 mM KOAc, 5 mM Mg(OAc)_2_, 1 mM TCEP) using a Biocomp Gradient Master and chilled at 4 °C overnight. Samples from cellular lysates were normalized by total protein with Pierce BCA assays (Thermofisher, 23225), and 200 µl lysate, ∼ 1mg of total protein, was loaded onto each gradient. Gradients were centrifuged at 40,000 rpm for 1:45 at 4 °C in open-top 12 ml PolyClear tubes using a Sw41 Ti swinging bucket rotor. Gradients were fractionated with a Biocomp piston fractionator and A_260_ profiles of the gradients were recorded with a Triax flow cell.

### Coomassie Staining and Autoradiography

Reactions to be analyzed by Coomassie stain and autoradiography were separated by gel electrophoresis using 4-20 % Tris Glycine TGX gels (Biorad, 5671095) run in 1 X Tris Glycine Running Buffer (Biorad, 1610732) at 150 V for 60 min. Afterwards, gels were fixed in Buffer X (40 % methanol, 10 % acetic acid), stained for 15 min in Coomassie R250 stain (0.1 % Coomassie R-250 (Thermofisher, 20278), 40 % methanol, 10 % acetic acid), and destained for 2 hours in Buffer V. Destained gels were imaged on a GelDoc Imager (Biorad, 12009077) before drying on a vacuum gel dryer (Biorad, Model 583) for 1 hour at 70 °C. Dried gels were exposed to magnetic phosphor screens (Cytiva, 28956476) for 30 min (substrate phosphorylation) or overnight (autophosphorylation) and imaged with a Typhoon Imager (Cytiva, 29187193) using 100 µM pixel size and 1000 V PMT.

### Immunoblotting

Samples from cellular lysates were normalized by total protein with Pierce BCA assays and 10 µg total protein was loaded in each gel lane. Kinase reactions were diluted 1:10 and 8 µl was loaded in each lane. Electrophoresis was performed as described in “Autoradiography.” Proteins were transferred to 0.2 µm Immun-Blot PVDF membranes (Biorad, 1620175) in 1X Trans-Blot Turbo Buffer (Biorad, 10026938) using a Trans-Blot Turbo Semi-Dry transfer system (Biorad, 1704150) with a 10 min, 2.5 A protocol.

Membranes were blocked with 5 % non-fat milk solids (Santa Cruz Biotechnology, sc-2324) in 1X TBST (20 mM Tris·HCl pH 7.5, 150 mM NaCl, 0.1 % Tween-20) for 1 hour. Primary antibodies were diluted in 5 % non-fat milk solids in 1X TBST and incubated with membranes overnight at 4 °C with gentle rocking. The next morning membranes were washed 4 times for 5 min in 1X TBST before addition of HRP-conjugated secondary antibodies diluted in 5 % non-fat milk solids in 1X TBST. Membranes were incubated with secondary antibodies for 1 hour at room temperature and then washed 4 times with 1X TBST for 5 min. Membranes were then dried, exposed for 2 min to SuperSignal Pico PLUS (Thermofisher, 34580) or Femto Maximum Sensitivity (Thermofisher, 34094) substrate, and imaged by chemiluminescence with a GelDoc Imager.

### Northern Blotting

Total RNA was extracted from cellular lysates by mixing 100 µl lysate with 1 ml TRIzol reagent (Thermofisher, 15596026). Samples were vortexed briefly and incubated 5 min at RT. 200 µl chloroform (Millipore Sigma, C2432) was added and the samples were mixed thoroughly and centrifuged at 12,000 x *g* for 15 min at 4 °C. The resulting aqueous phase was mixed with 500 µl ice-cold isopropanol, incubated for 10 min at 4 °C, and centrifuged at 12,000 x *g* for 10 min at 4 °C. The precipitated RNA pellet was washed twice with 75 % ethanol, air dried, and resuspended in 20 µl TE buffer (Thermofisher, AM9849). RNA concentration was measured by Qubit Broad Range (BR) assay (Thermofisher, Q10210) and 10 µg was mixed with equal volume 2X RNA loading dye (NEB, B0363S), heated at 95 °C for 3 min, cooled on ice, and loaded on a 10 % TBE-Urea PAGE gel (Biorad, 3450089). Gels were run at 200 V for 90 min in 1X TBE buffer (Thermofisher, 15581044) and transferred to a Hybond-N+ membrane (Cytiva, 15581044) in 1X TBE using a TransBlot Turbo Semi-Dry transfer system in 1X TBE with a 30 min 1.0 A protocol. Membranes were crosslinked using a Stratalinker 2400 UV crosslinker (Agilent Technologies, 400075) using the “Auto” setting and then washed twice with 2X SSC buffer (Thermofisher, AM9763).

Prior to hybridization, membranes were incubated in NorthernMax hybridization buffer (Thermofisher, AM8677) at 42 °C for 60 min with gentle rotation. 50 pmol probe was labeled with 50 pmol of γ^32^P-labeled ATP (Revvity, BLU035C005MC) using T4 polynucleotide kinase (NEB, M0201S) in 50 µl reactions. Excess nucleotides were removed with MicroSpin G-50 columns (Cytiva, 27533001). 10 µl Phosphorylated probe was diluted in 25 ml pre-warmed hybridization buffer and incubated with membranes at 42 °C overnight with gentle rotation. The next morning the membranes were washed 3 times for 20 min with 2X SSC for 20 before exposure to a magnetic phosphor screen and imaging by autoradiography with a Typhoon imager.

### Data Analysis

Autoradiographs and Coomassie stained images were analyzed using ImageJ (https://imagej.net/ij/). ROIs were contrast adjusted using the Adjust Brightness and Contrast “Auto” setting and background corrected with a 50 pixel rolling ball radius. Band intensities were extracted using the built in “Analyze Gel” toolkit. The resulting values were normalized in Matlab (www.mathworks.com/). For Coomassie stained gels, the integrated intensity of the upper, phosphorylated, eIF2α-NTD band was divided by the sum of the upper and lower bands to yield the fraction phosphorylated. For each kinetic experiment, the fraction phosphorylated measured by Coomassie stain at the longest timepoint was used to normalize the integrated intensity of the autoradiographic signals of substrate phosphorylation to produce the fraction phosphorylated values reported. The autoradiographic signals for autophosphorylation lack a Coomassie stain normalization and were instead normalized to the highest single band intensity on the gel to produce a relative phosphorylation. The fractional phosphorylations from substrate phosphorylation time courses were plotted, along with standard error of the mean (SEM) at each timepoint, in GraphPad Prism (https://www.graphpad.com/) and fit with a two-term hypo-exponential model:

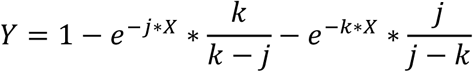

The relative phosphorylations from autophosphorylation were plotted in Graphpad, along with SEM at each timepoint and fit with the software’s default exponential association model. Fits were differentiated with the GraphPad “Analysis, Differentiate” function with no smoothing, and the maximum of the derivative was plotted as a function of concentration to establish the dosage response of agonism or antagonism. These curves were fit with GraphPad’s 3 or 4 parameter dosage responses curves or the Michaelis-Menten equation using default settings to yield estimates of EC_50_ or IC_50_.

### Structural Modeling

Structural predictions were generated with Alphafold3 (*79*) in the Google Collab environment (https://alphafoldserver.com/) using default settings. The residues and nucleotides included in each model are indicated in the figure legends. Structural predictions were visualized in UCSF ChimeraX (https://www.cgl.ucsf.edu/chimerax/).

### Sequence Alignment

GCN2 protein sequences from all *Chordata* species annotated were downloaded from the NCBI RefSeq database (https://www.ncbi.nlm.nih.gov/refseq/). Sequences were limited to one, longest, isoform per species. Sequences were aligned using Clustal Omega (*80*) in JalView (https://www.jalview.org/).

## Supplementary Figures

**Figure S1.**
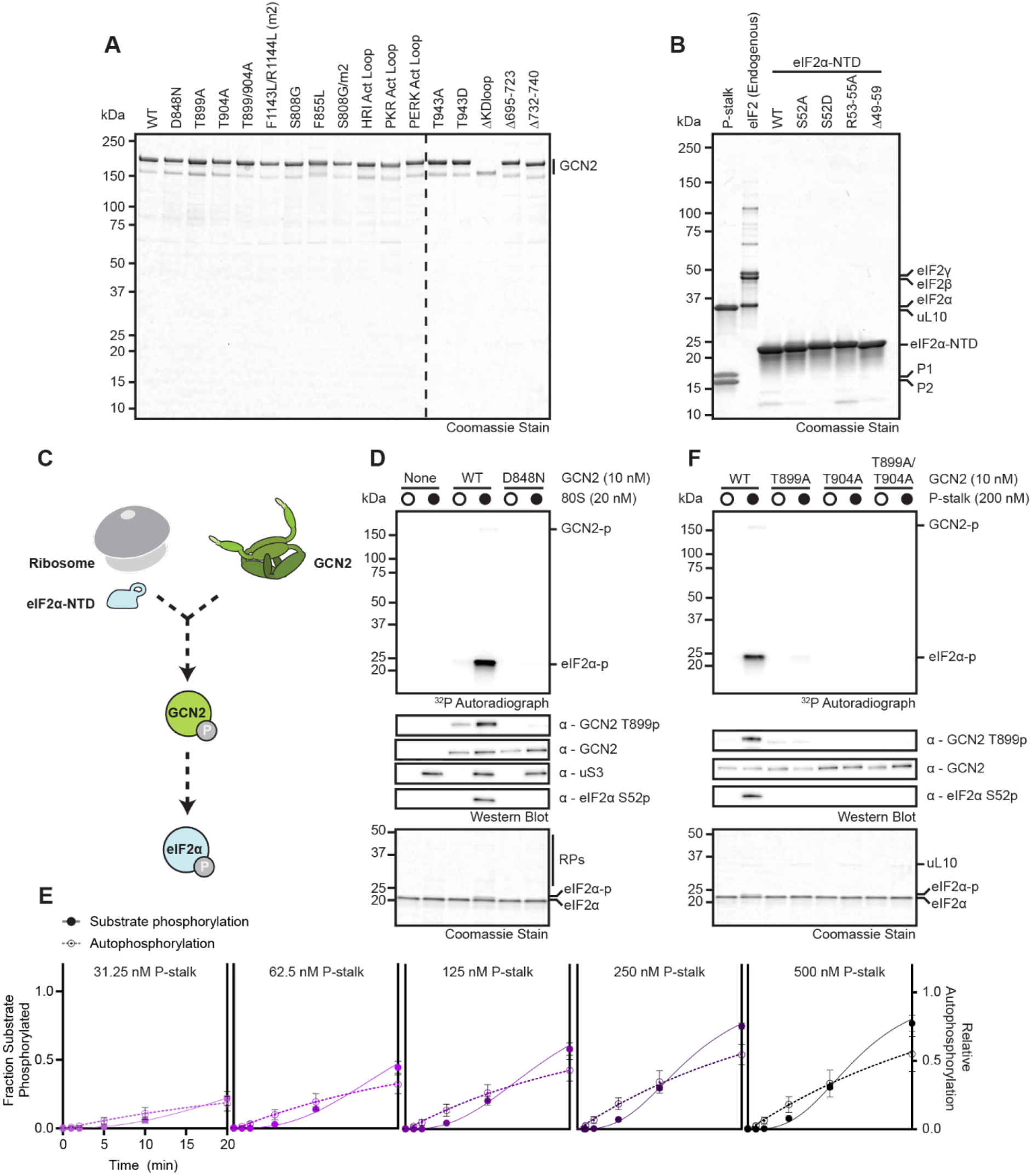
**(A)** Coomassie stained gel with 1-2 pmol of recombinant GCN2 variants used in this study. **(B)** Coomassie stained gel with 35 pmol purified P-stalk, 10 pmol eIF2 trimer, and 75 pmol of all eIF2α-NTD variants used in this study. **(C)** Schematic of a kinase assay using recombinant GCN2 (green), 80S ribosomes (gray), and eIF2α-NTD (light blue). **(D)** Coomassie stain, autoradiographs, and immunoblots with the indicated antibodies of kinase assays as in (B) in the presence or absence of 20 nM 80S ribosomes with 10 nM wild-type (WT) or kinase-dead D848N GCN2 and 2 µM eIF2α-NTD. **(E)** Intensities of autophosphorylation and substrate phosphorylation autoradiographs at the indicated P-stalk concentrations as in (Figure 1D) plotted and fit with single or hypo-exponential functions, respectively. Error bars represent S.E.M. of 4 independent replicates. **(F)** Coomassie stain, autoradiograph, and immunoblot with the indicated antibodies of kinase assays as in (Figure 1A) in the presence or absence of 200 nM P-stalk with 10 nM of the indicated GCN2 variants and 2 µM eIF2α-NTD.

**Figure S2.**
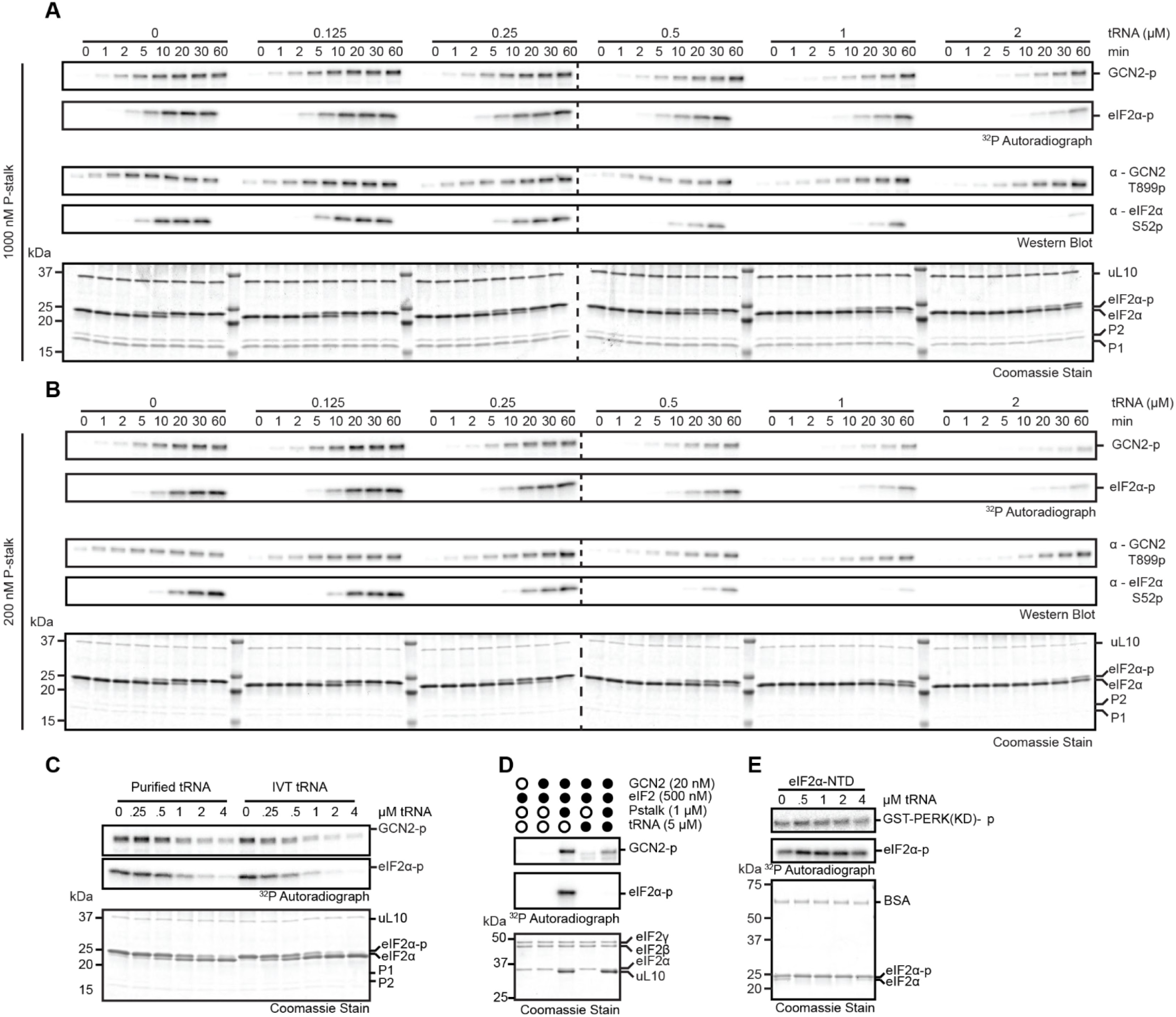
**(A)** Coomassie stain, autoradiographs, and immunoblots with the indicated antibodies of kinetic kinase assays with 1000 nM P-stalk and a titration of tRNA as outlined in (Figure 2D). **(B)** Coomassie stain, autoradiographs, and immunoblots with the indicated antibodies of kinetic kinase assays with 200 nM P-stalk and a titration of tRNA as outlined in (Figure 2D). **(C)** Coomassie stain and autoradiographs of kinase assays with 20 nM GCN2, 200 nM P-stalk, 2 µM eIF2α-NTD, and a titration of purified, deacylated pig liver tRNA or *in vitro* transcribed Leu_CAG_ tRNA. **(D)** Coomassie stain and autoradiographs of kinase assays with 20 nM GCN2, 200 nM P-stalk, 500 nM purified endogenous eIF2 trimer, and 5 µM tRNA as indicated. **(E)** Coomassie stain and autoradiographs of kinase assays with 20 nM GST-PERK kinase domain (KD), 2 µM of eIF2α-NTD, and a titration of tRNA.

**Figure S3.**
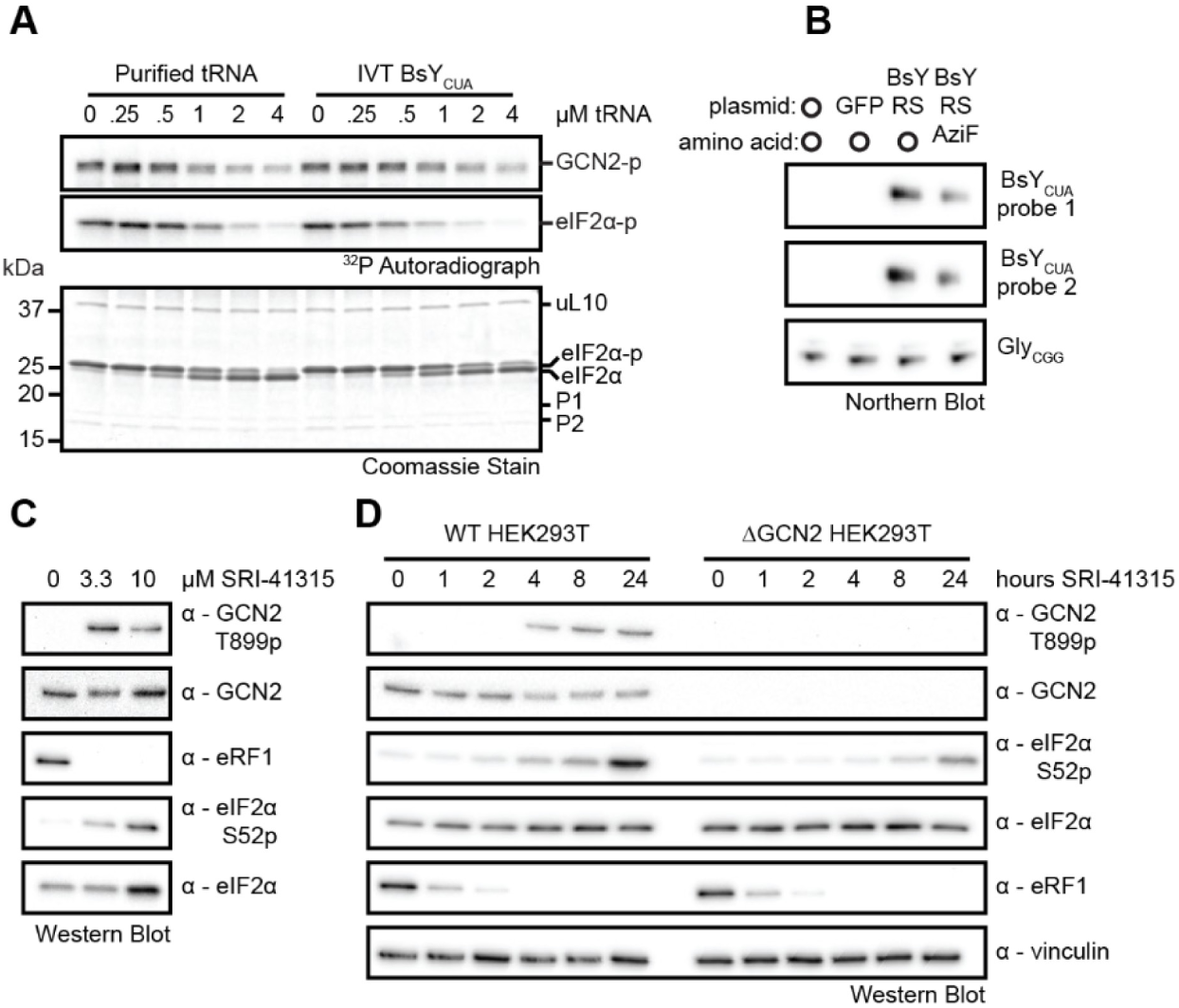
**(A)** Coomassie stain and autoradiographs of kinase assays with 20 nM GCN2, 200 nM P-stalk, 2 µM eIF2α-NTD, and a titration of purified, deacylated pig liver tRNA or *in vitro* transcribed BsY_CUA_ amber suppression tRNA. **(B)** Northern blots with the indicated probes of BsY_CUA_ and Gly_GCC_ expression in cells transfected with no DNA, GFP (pFD042), or BsY amber suppression plasmids (pIRE4-Azi) in the presence or absence of Azido-Phenylalanine (AzF). **(C)** Immunoblots of GCN2 activity in HEK293T cells treated with 0, 3.3, and 10 µM SRI-41315 for 4 h to induce degradation of eRF1. **(D)** Immunoblots of GCN2 activity in wild-type (WT) and GCN2 KO HEK293T cells treated with 10 µM SRI-41315 for the indicated time course to induce degradation of eRF1.

**Figure S4.**
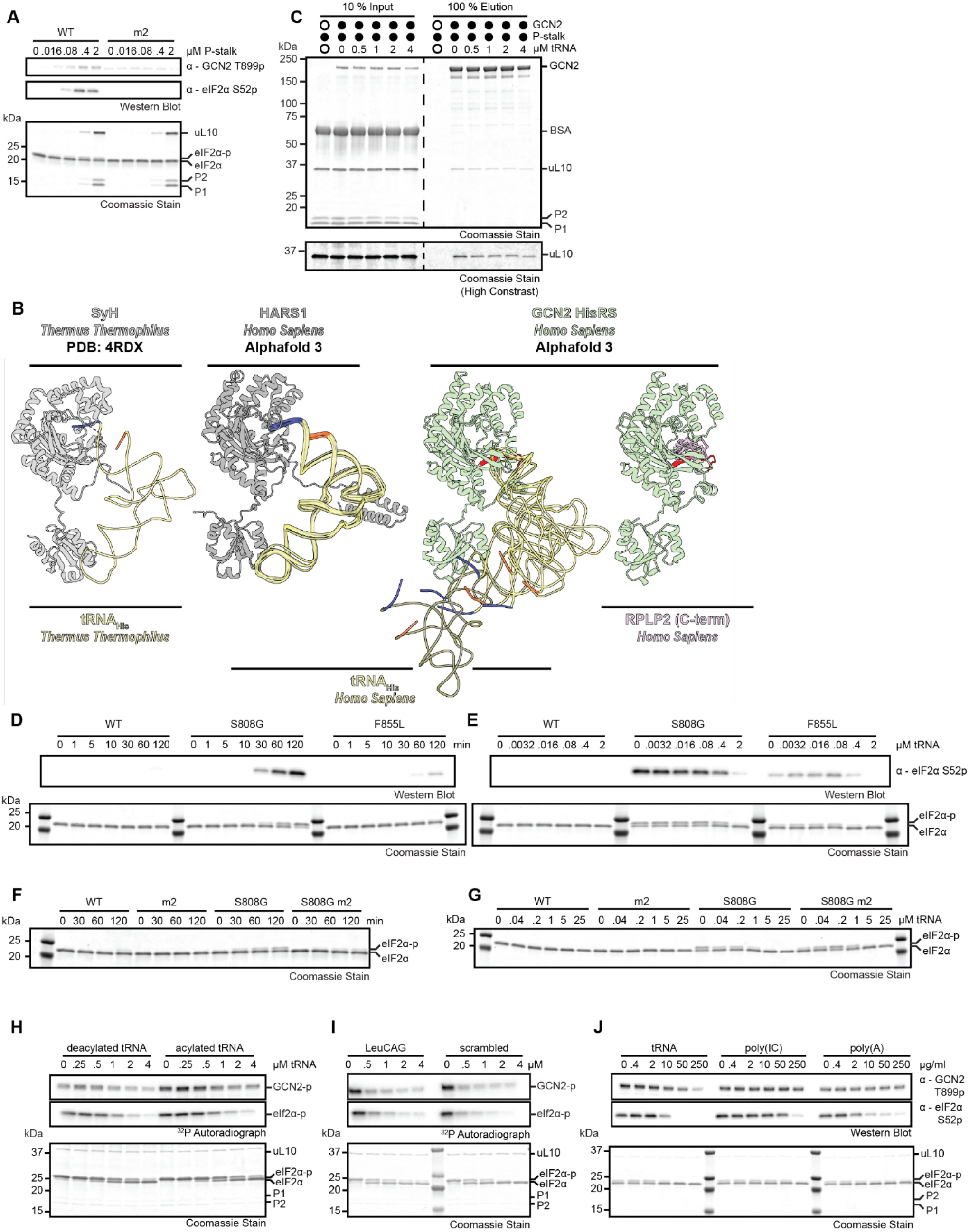
**(A)** Coomassie stain and immunoblots with the indicated antibodies of kinase assays with 20 nM wild-type (WT) or F1143L/R1144L (m2) GCN2, 2 µM eIF2α-NTD, and the indicated titration of P-stalk. **(B)** Coomassie stains of co-immunoprecipitation experiments with 200 nM Strep-tagged GCN2 and 1 µM P-stalk in the presence of a titration of tRNA. The region including uL10 is displayed at two separate contrast levels for clarity. **(C)** Structural models of *Thermus thermophilus* Histidine tRNA Synthetase (SyH; residues 2-421, *tt*SyH; gray) with tRNA_His_ (*tt*tRNA-His-GTG-1-1; yellow) (Crystal structure, PDB 4RDX (*81*)), *Homo sapiens* Histidine tRNA Synthetase (HARS, residues 1-509; *hs*HARS1, gray) with tRNA_His_ (*hs*tRNA-His-GTG-1-1; yellow) (Alphafold3, top 5 predictions), *Homo sapiens* GCN2 Histidine tRNA Synthetase-like domain (HisRS; residues 1022-1493, *hs*EIF2AK4; green) with tRNA_His_ (*hs*tRNA-His-GTG-1-1; yellow) (Alphafold3, top 5 predictions), and *Homo sapiens* GCN2 HisRS (HisRS; residues 1022-1493, *hs*EIF2AK4; green) with the C-terminal tail of P-stalk protein 2 (RPLP2; residues 106-115, *hs*RLA2; pink) (Alphafold3, top 5 predictions). The 5’ and 3’ ends of the tRNAs are colored orange and blue, respectively. The motif 2 (m2) loop of the GCN2 HisRS domain is colored red. **(D)** Coomassie stain and immunoblot with the indicated antibody of kinase assays with 20 nM of the indicated GCN2 variants and 2 µM eIF2α-NTD at the indicated time points. **(E)** As in (D), but at 60 min only with the indicated titration of tRNA. **(F)** Coomassie stain as in (D) with the indicated GCN2 variants. **(G)** As in (E) but at 60 min only with the indicated titration of tRNA. **(H)** Coomassie stain and autoradiographs of kinase assays with 20 nM GCN2, 200 nM P-stalk, 2 µM eIF2α-NTD, and a titration of partially acylated and deacylated pig liver tRNA. **(I)** As in (H) but with titrations of *in vitro* transcribed Leu_CAG_ or scrambled tRNAs. **(J)** Coomassie stain and immunoblots with the indicated antibodies of kinase assays as in (H), but with titrations of equivalent masses of purified tRNA, polyIC, and polyA.

**Figure S5.**
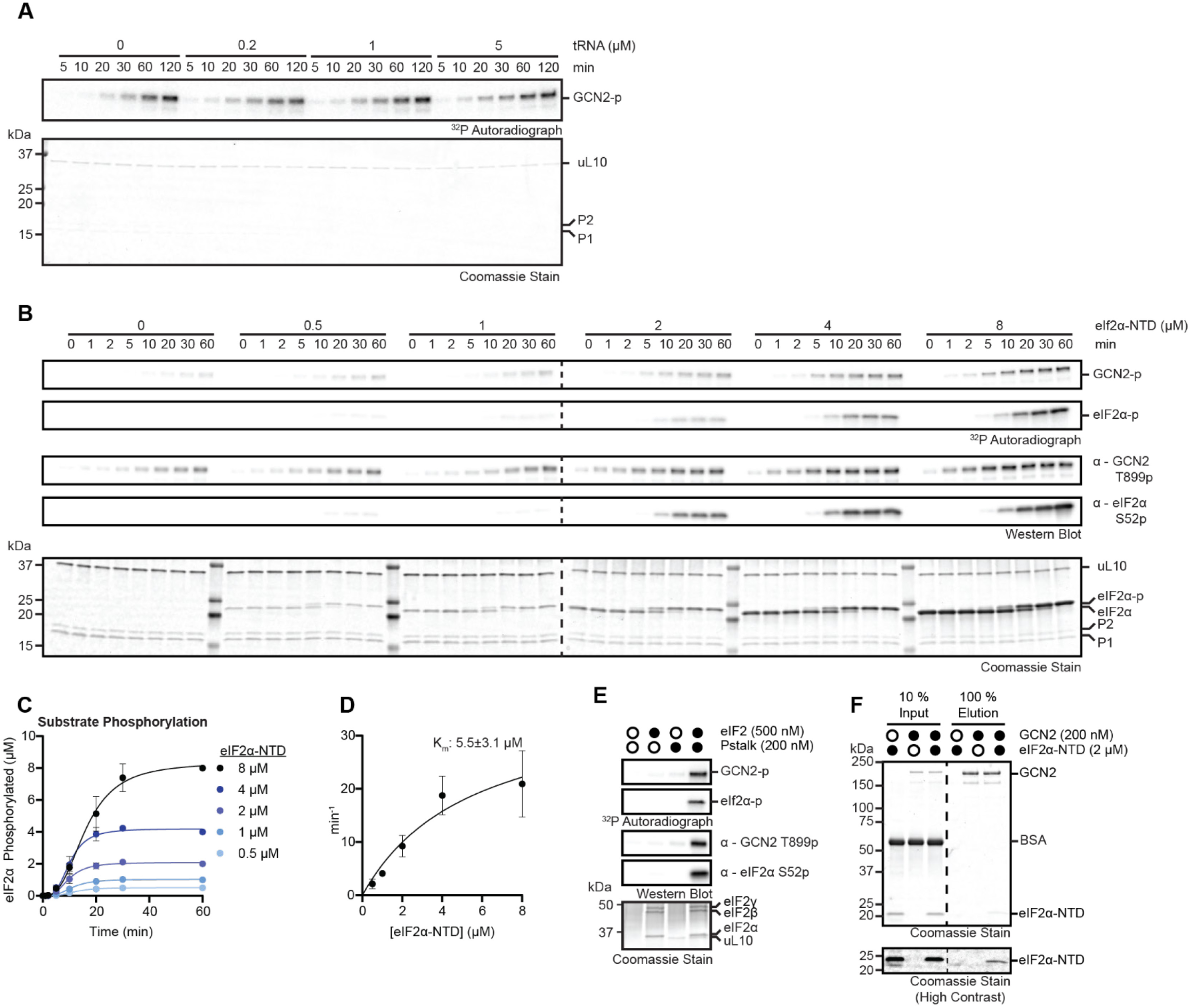
**(A)** Coomassie stain and autoradiographs of kinetic kinase assays with 200 nM P-stalk and a titration of tRNA as outlined in (Figure 4B). **(B)** Coomassie stain, autoradiographs, and immunoblots with the indicated antibodies of kinetic kinase assays with 1000 nM P-stalk and a titration of eIF2α-NTD as outlined in (Figure 4G). **(C)** Intensities of substrate autoradiographs as in (B) plotted and fit with hypo-exponential functions. Error bars represent S.E.M. of 2 independent replicates. **(D)** V_0_ in (C) as a function eIF2α-NTD concentration fit with the Michaelis-Menten equation (see Methods). Error bars represent S.E.M. of 2 different replicates. Error on reported K_m_ is the standard error of the fit. **(E)** Coomassie stain, autoradiographs, and immunoblots with the indicated antibodies of 30 min kinase assays with 20 nM GCN2 in the presence or absence of 200 nM P-stalk and 500 nM purified endogenous eIF2 trimer. **(F)** Coomassie stains from co-immunoprecipitation experiments with 200 nM Strep-tagged GCN2 and 2 µM eIF2α-NTD. The region including eIF2α-NTD is displayed at two separate contrast levels for clarity.

**Figure S6.**
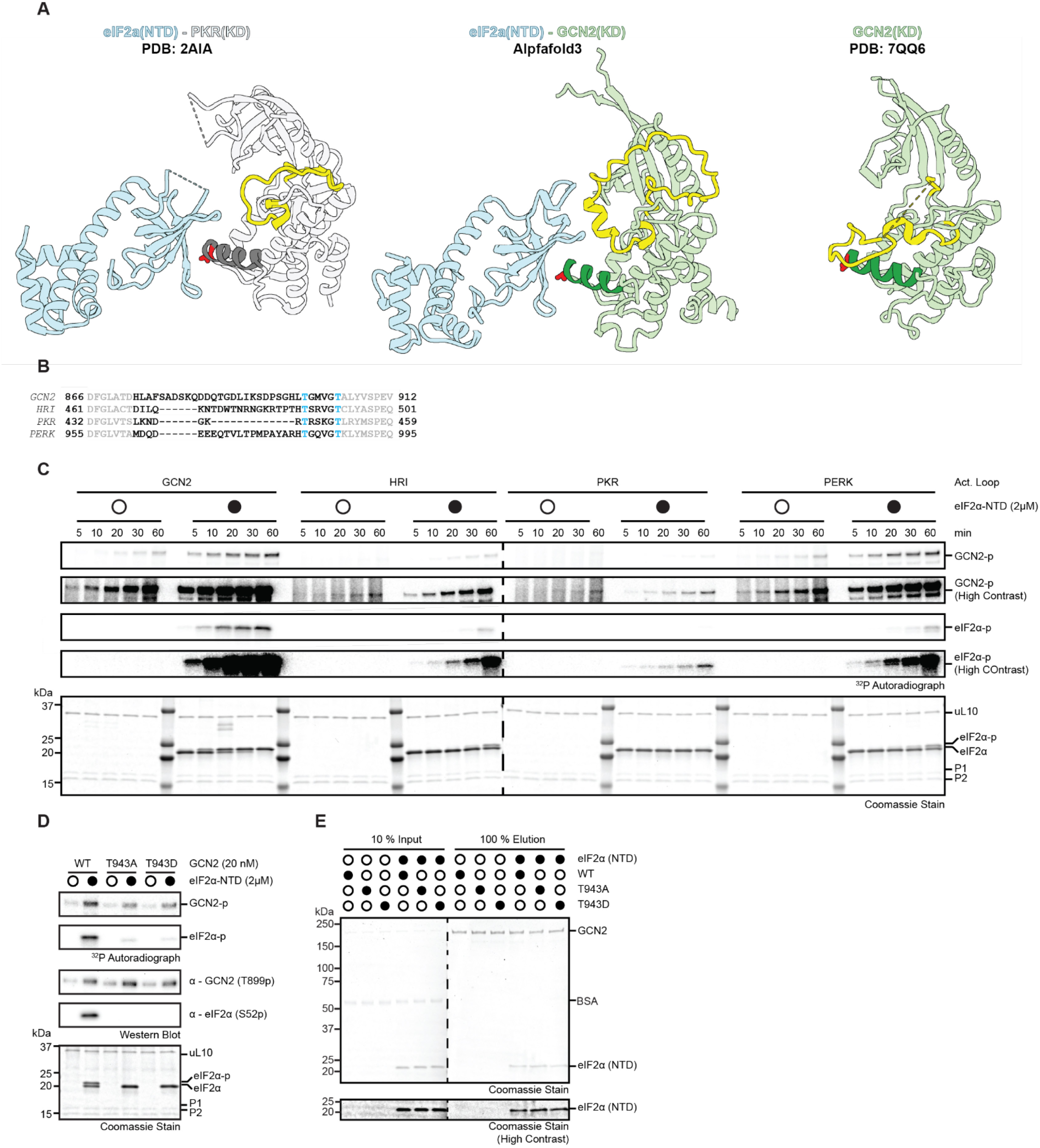
**(A)** Structural models of *Homo sapiens* PKR kinase domain (PKR(KD); residues 258-541, *hs*EIF2AK2, grey) with *Saccharomyces cerevisiae* eIF2α-NTD (eIF2α-NTD; residues 5-176, *sc*SUI2; light blue) (Crystal structure, PDB 2A1A (*38*)), *Homo sapiens* GCN2 kinase domain (GCN2(KD); residues 585-1016, *hs*EIF2AK4; green) with *Homo sapiens* eIF2α-NTD (eIF2α-NTD; residues 2-185, *hs*IF2A; light blue) (Alphafold3), *Homo sapiens* GCN2 kinase domain (GCN2(KD); residues 584-1006, *hs*EIF2AK4; green) (Crystal structure, PDB 7QQ6(*15*)). The αG helices of the kinase domains are colored in darker shades, with the Threonine residues corresponding to T943 in red. The activation loops of the kinases are colored yellow. **(B)** Clustal Omega alignments of the activation segments of the human ISR kinases. The disorded activation loop is indicated in dark font. The phosphorylated Threonine residues are colored blue. **(C)** Coomassie stains and autoradiographs from kinetic kinase assays with 20 nM of chimeric GCN2 constructs whose activation loop has been replaced by that of HRI, PKR, or PERK, with 200 nM P-stalk, in the presence or absence of 2 µM eIF2α-NTD. Autoradiographs are displayed at two separate contrast levels for clarity. **(D)** Coomassie stain, autoradiographs, and immunoblots with the indicated antibodies of 10 min kinase assays with 20 nM of the indicated GCN2 variants and 200 nM P-stalk in the presence or absence of 2 µM eIF2α-NTD. **(E)** Coomassie stains from co-immunoprecipitation experiments with 200 nM Strep-tagged GCN2 variants and 2 µM eIF2α-NTD. The region including eIF2α-NTD is displayed at two separate contrast levels for clarity.

**Figure S7.**
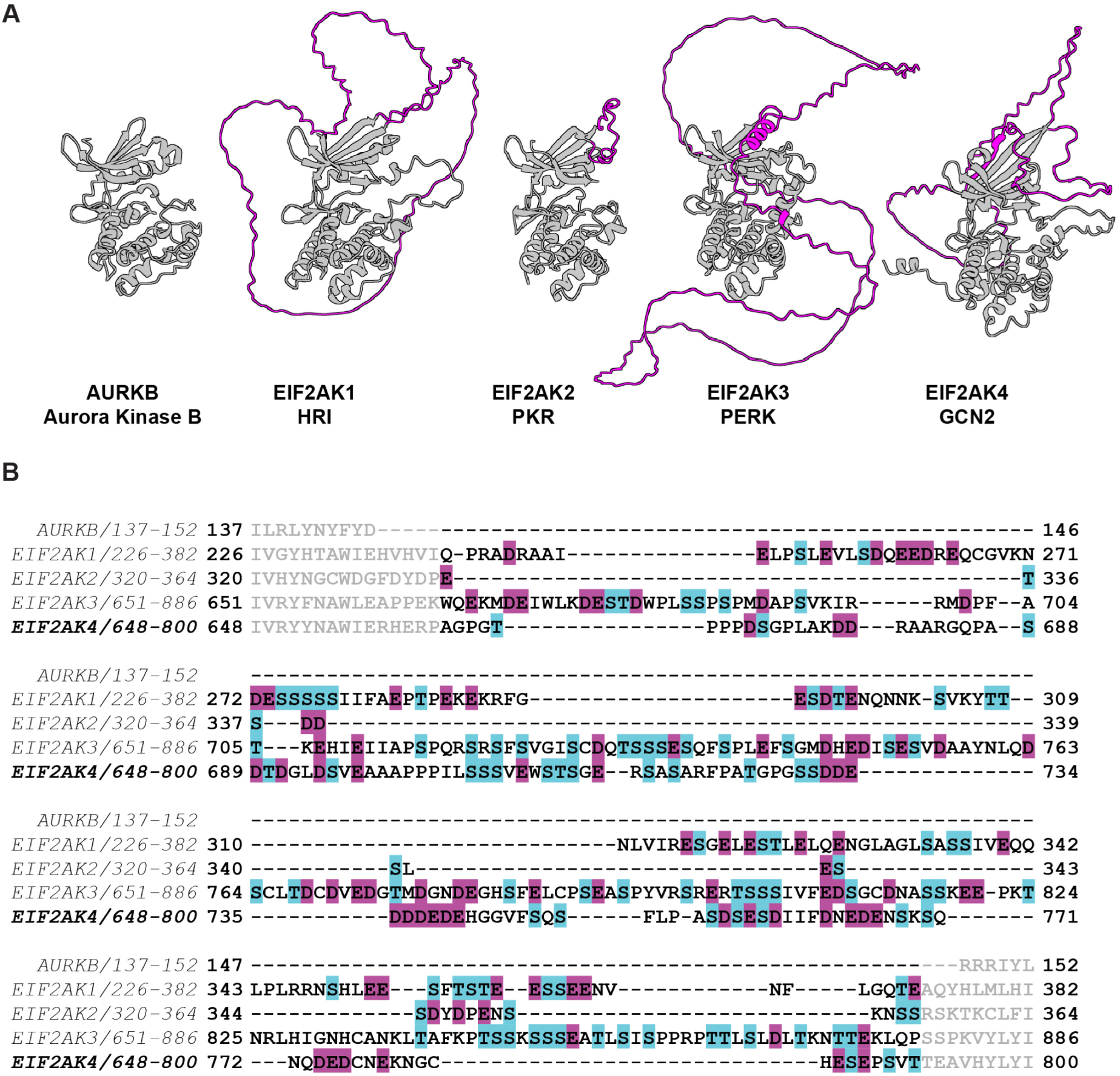
**(A)** Alphafold3 structural models of *Homo sapiens* Aurora Kinase B kinase domain (AURKB; residues 77-327, *hs*AURKB), *Homo sapiens* HRI kinase domain (HRI; residues 167-583. *hs*EIF2AK1), *Homo sapiens* PKR kinase domain (PKR; residues 267-538, *hs*EIF2AK2), *Homo* sapiens PERK kinase domain (PERK; residues 593-1077, *hs*EIF2AK3), Homo *sapiens* GCN2 kinase domains (GCN2; residues 585-1016, *hs*EIF2AK4). The kinase domain insertion loop is colored in magenta. **(B)** Clustal Omega alignment of the insertion loops of the kinases in (A). The insertion loop is indicated by darker font. S/T residues are colored in cyan. D/E residues are colored in magenta.

**Figure S8.**
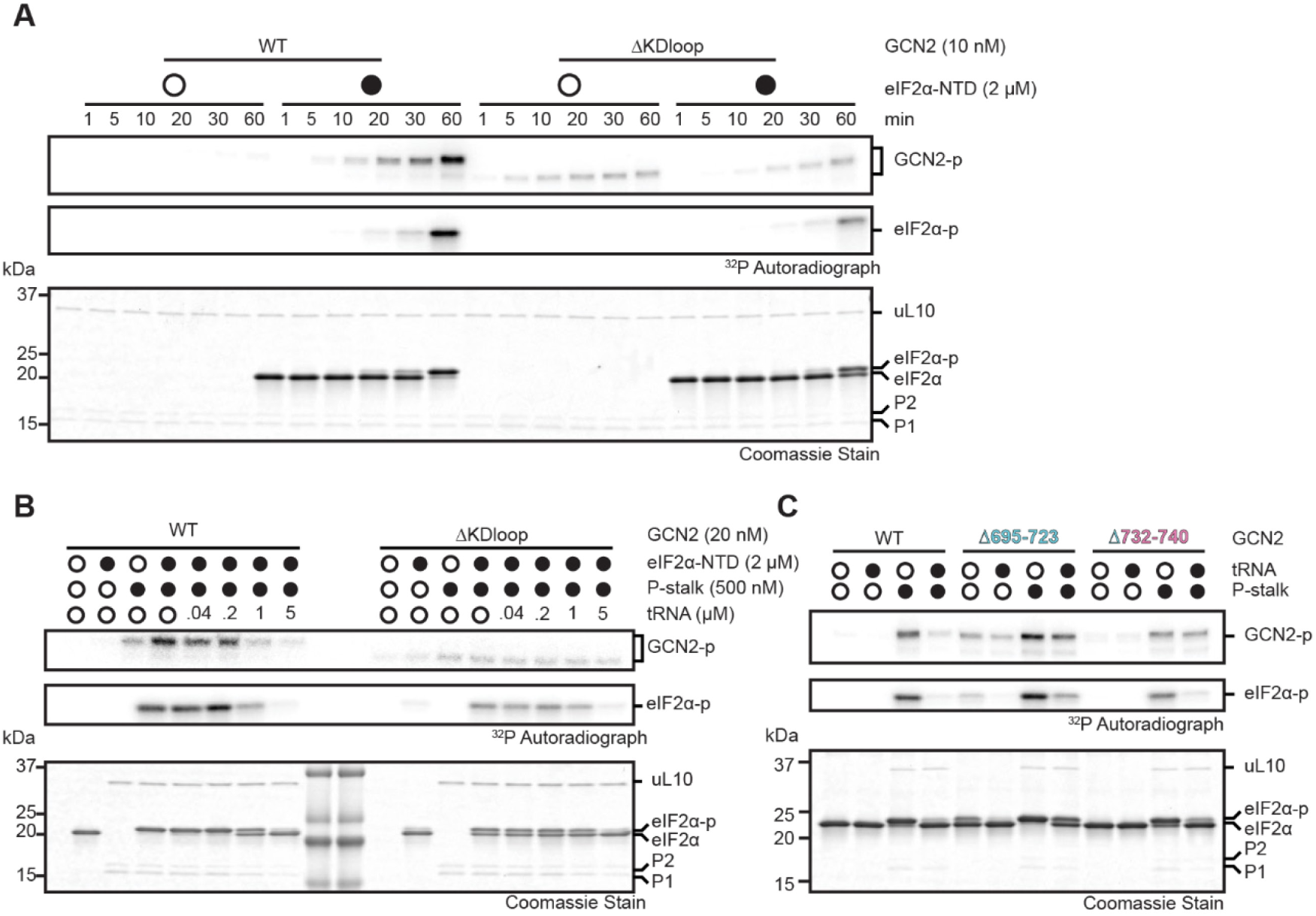
**(A)** Coomassie stain and autoradiographs of kinetic kinase assays with 10 nM wild-type (WT) or kinase insertion loop deletion (ΔKDloop) GCN2 and 200 nM P-stalk in the presence or absence of 2 µM eIF2α-NTD. **(B)** Coomassie stain and autoradiographs of 30 min kinase assays with 20 nM wild-type (WT) or kinase insertion loop deletion (ΔKDloop) GCN2, 500 nM P-stalk, 2 µM eIF2α-NTD, and the indicated tRNA titration. **(C)** Coomassie stain and autoradiographs of 30 min kinase assays with the indicated GCN2 variants and 2 µM eIF2α-NTD in the presence or absence of 200 nM P-stalk and 5 µM tRNA.

**Figure S9.**
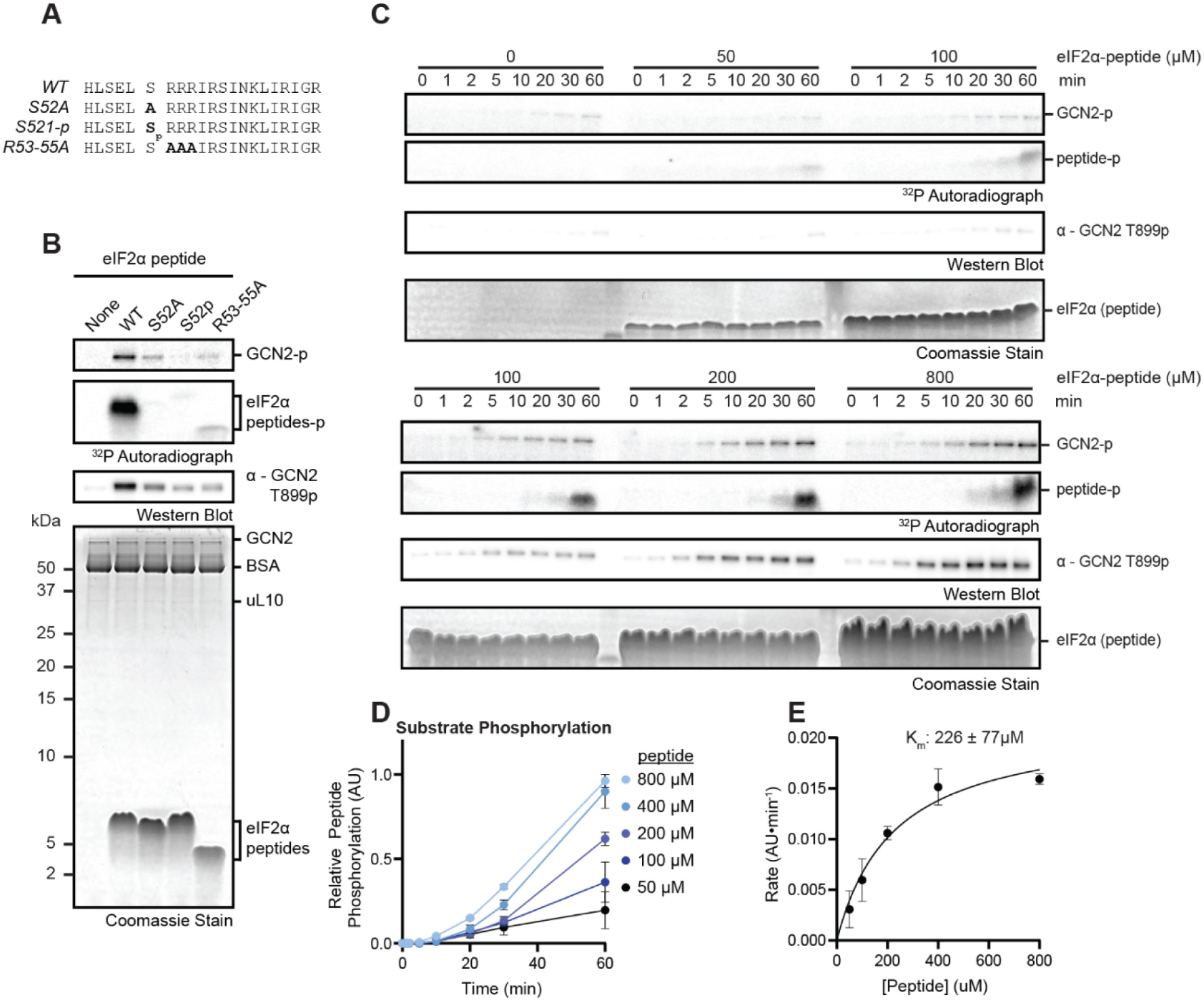
**(A)** eIF2α phosphorylated loop peptides. Mutated residues are indicated by darker font. **(B)** Coomassie stain, autoradiographs, and immunoblot with the indicated antibody of 30 min kinase assays with 20 nM GCN2, 200 nM P-stalk, and the indicated eIF2α peptides. **(C)** Coomassie stain, autoradiographs, and immunoblots with the indicated antibody of kinetic kinase assays with a titration of P-stalk as outlined in (Figure 6E). **(D)** Intensities of substrate autoradiographs as in (B) plotted and fit with hypo-exponential functions. Error bars represent S.E.M. of 2 independent replicates. **(E)** V_0_ in (D) as a function eIF2α peptide concentration fit with the Michaelis-Menten equation (see Methods). Error bars represent S.E.M. of 2 different replicates. Error on reported K_m_ is the standard error of the fit.

## References

1. T. E. Dever, J. J. Chen, G. N. Barber, A. M. Cigan, L. Feng, T. F. Donahue, I. M. London, M. G. Katze, A. G. Hinnebusch, Mammalian eukaryotic initiation factor 2 alpha kinases functionally substitute for GCN2 protein kinase in the GCN4 translational control mechanism of yeast. Proc. Natl. Acad. Sci. U. S. A. 90, 4616–4620 (1993).

2. H. P. Harding, I. Novoa, Y. Zhang, H. Zeng, R. Wek, M. Schapira, D. Ron, Regulated translation initiation controls stress-induced gene expression in mammalian cells. Mol. Cell 6, 1099–1108 (2000).

3. H. P. Harding, Y. Zhang, H. Zeng, I. Novoa, P. D. Lu, M. Calfon, N. Sadri, C. Yun, B. Popko, R. Paules, D. F. Stojdl, J. C. Bell, T. Hettmann, J. M. Leiden, D. Ron, An integrated stress response regulates amino acid metabolism and resistance to oxidative stress. Mol. Cell 11, 619–633 (2003).

4. A. G. Hinnebusch, G. R. Fink, Positive regulation in the general amino acid control of Saccharomyces cerevisiae. Proc. Natl. Acad. Sci. U. S. A. 80, 5374–5378 (1983).

5. M. D. Penn, B. Galgoci, H. Greer, Identification of AAS genes and their regulatory role in general control of amino acid biosynthesis in yeast. Proc. Natl. Acad. Sci. U. S. A. 80, 2704–2708 (1983).

6. J. Deng, H. P. Harding, B. Raught, A.-C. Gingras, J. J. Berlanga, D. Scheuner, R. J. Kaufman, D. Ron, N. Sonenberg, Activation of GCN2 in UV-irradiated cells inhibits translation. Curr. Biol. CB 12, 1279–1286 (2002).

7. R. Ishimura, G. Nagy, I. Dotu, J. H. Chuang, S. L. Ackerman, Activation of GCN2 kinase by ribosome stalling links translation elongation with translation initiation. eLife 5, e14295 (2016).

8. C. C.-C. Wu, A. Peterson, B. Zinshteyn, S. Regot, R. Green, Ribosome Collisions Trigger General Stress Responses to Regulate Cell Fate. Cell 182, 404–416.e14 (2020).

9. M. J. Derisbourg, M. D. Hartman, M. S. Denzel, Perspective: Modulating the integrated stress response to slow aging and ameliorate age-related pathology. Nat. Aging 1, 760–768 (2021).

10. L. T. Gold, G. R. Masson, GCN2: roles in tumour development and progression. Biochem. Soc. Trans. 50, 737–745 (2022).

11. N. Nameki, M. Yoneyama, S. Koshiba, N. Tochio, M. Inoue, E. Seki, T. Matsuda, Y. Tomo, T. Harada, K. Saito, N. Kobayashi, T. Yabuki, M. Aoki, E. Nunokawa, N. Matsuda, N. Sakagami, T. Terada, M. Shirouzu, M. Yoshida, H. Hirota, T. Osanai, A. Tanaka, T. Arakawa, P. Carninci, J. Kawai, Y. Hayashizaki, K. Kinoshita, P. Güntert, T. Kigawa, S. Yokoyama, Solution structure of the RWD domain of the mouse GCN2 protein. Protein Sci. Publ. Protein Soc. 13, 2089–2100 (2004).

12. Z. Hei, S. Wu, L. Zheng, J. Zhou, Z. Liu, J. Wang, P. Fang, Crystal structures reveal a novel dimer of the RWD domain of human general control nonderepressible 2. Biochem. Biophys. Res. Commun. 549, 164–170 (2021).

13. Y. Liu, J. Misra, D. R. Bhowmik, B. O’Boyle, K. A. Staschke, N. Kannan, R. C. Wek, N. Jura, Structural basis for pseudokinase-mediated regulation of GCN2 in the integrated stress response. Proc. Natl. Acad. Sci. U. S. A. 123, e2526598123 (2026).

14. A. K. Padyana, H. Qiu, A. Roll-Mecak, A. G. Hinnebusch, S. K. Burley, Structural basis for autoinhibition and mutational activation of eukaryotic initiation factor 2alpha protein kinase GCN2. J. Biol. Chem. 280, 29289–29299 (2005).

15. T. Maia de Oliveira, V. Korboukh, S. Caswell, J. J. Winter Holt, M. Lamb, A. W. Hird, R. Overman, The structure of human GCN2 reveals a parallel, back-to-back kinase dimer with a plastic DFG activation loop motif. Biochem. J. 477, 275–284 (2020).

16. K. Solorio-Kirpichyan, X. Fan, D. Golovenko, A. A. Korostelev, N. Yan, A. Korennykh, Cryo-EM structure of histidyl-tRNA synthetase-like domain reveals activating crossed helices at the core of GCN2. PNAS Nexus 3, pgae528 (2024).

17. C. Bou-Nader, S. Gaikwad, S. Bahmanjah, F. Zhang, A. G. Hinnebusch, J. Zhang, Gcn2 structurally mimics and functionally repurposes the HisRS enzyme for the integrated stress response. Proc. Natl. Acad. Sci. U. S. A. 121, e2409628121 (2024).

18. J. Z. Yin, A. F. A. Keszei, S. Houliston, F. Filandr, J. Beenstock, S. Daou, J. Kitaygorodsky, D. C. Schriemer, M. T. Mazhab-Jafari, A.-C. Gingras, F. Sicheri, The HisRS-like domain of GCN2 is a pseudoenzyme that can bind uncharged tRNA. Struct. Lond. Engl. 1993 32, 795–811.e6 (2024).

19. H. He, I. Singh, S. A. Wek, S. Dey, T. D. Baird, R. C. Wek, M. M. Georgiadis, Crystal structures of GCN2 protein kinase C-terminal domains suggest regulatory differences in yeast and mammals. J. Biol. Chem. 289, 15023–15034 (2014).

20. R. C. Wek, B. M. Jackson, A. G. Hinnebusch, Juxtaposition of domains homologous to protein kinases and histidyl-tRNA synthetases in GCN2 protein suggests a mechanism for coupling GCN4 expression to amino acid availability. Proc. Natl. Acad. Sci. U. S. A. 86, 4579–4583 (1989).

21. S. A. Wek, S. Zhu, R. C. Wek, The histidyl-tRNA synthetase-related sequence in the eIF-2 alpha protein kinase GCN2 interacts with tRNA and is required for activation in response to starvation for different amino acids. Mol. Cell. Biol. 15, 4497–4506 (1995).

22. S. Zhu, A. Y. Sobolev, R. C. Wek, Histidyl-tRNA synthetase-related sequences in GCN2 protein kinase regulate in vitro phosphorylation of eIF-2. J. Biol. Chem. 271, 24989–24994 (1996).

23. J. Dong, H. Qiu, M. Garcia-Barrio, J. Anderson, A. G. Hinnebusch, Uncharged tRNA activates GCN2 by displacing the protein kinase moiety from a bipartite tRNA-binding domain. Mol. Cell 6, 269–279 (2000).

24. R. Gupta, A. G. Hinnebusch, Differential requirements for P stalk components in activating yeast protein kinase Gcn2 by stalled ribosomes during stress. Proc. Natl. Acad. Sci. U. S. A. 120, e2300521120 (2023).

25. K. Worner, K. R. Maschhoff, G. M. Schuh, W. Hu, GCN2 monitors mRNA translation termination. Mol. Cell 85, 4575–4586.e5 (2025).

26. C. Zhou, M. Zhang, J. Murray, J. Paulo, S. Gygi, S. Shao, M. Whitman, T. Keller, GCN1 couples GCN2 to ribosomal state to initiate amino acid response pathway signaling. Science 390, eads8728 (2025).

27. A. A. Pochopien, B. Beckert, S. Kasvandik, O. Berninghausen, R. Beckmann, T. Tenson, D. N. Wilson, Structure of Gcn1 bound to stalled and colliding 80S ribosomes. Proc. Natl. Acad. Sci. U. S. A. 118, e2022756118 (2021).

28. A. Jiménez-Díaz, M. Remacha, J. P. G. Ballesta, J. J. Berlanga, Phosphorylation of initiation factor eIF2 in response to stress conditions is mediated by acidic ribosomal P1/P2 proteins in Saccharomyces cerevisiae. PloS One 8, e84219 (2013).

29. A. J. Inglis, G. R. Masson, S. Shao, O. Perisic, S. H. McLaughlin, R. S. Hegde, R. L. Williams, Activation of GCN2 by the ribosomal P-stalk. Proc. Natl. Acad. Sci. U. S. A. 116, 4946–4954 (2019).

30. H. P. Harding, A. Ordonez, F. Allen, L. Parts, A. J. Inglis, R. L. Williams, D. Ron, The ribosomal P-stalk couples amino acid starvation to GCN2 activation in mammalian cells. eLife 8, e50149 (2019).

31. J. Misra, K. R. Carlson, D. F. Spandau, R. C. Wek, Multiple mechanisms activate GCN2 eIF2 kinase in response to diverse stress conditions. Nucleic Acids Res. 52, 1830–1846 (2024).

32. T. E. Dever, L. Feng, R. C. Wek, A. M. Cigan, T. F. Donahue, A. G. Hinnebusch, Phosphorylation of initiation factor 2 alpha by protein kinase GCN2 mediates gene-specific translational control of GCN4 in yeast. Cell 68, 585–596 (1992).

33. R. Gupta, S. Gaikwad, H. Qui, C. Bou-Nader, J. Zhang, A. G. Hinnebusch, Purification and Analysis of eIF2α Phosphorylation by Stress-Activated Protein Kinase Gcn2 from S. cerevisiae. Methods Mol. Biol. 2882, 195–220 (2025).

34. P. R. Romano, M. T. Garcia-Barrio, X. Zhang, Q. Wang, D. R. Taylor, F. Zhang, C. Herring, M. B. Mathews, J. Qin, A. G. Hinnebusch, Autophosphorylation in the Activation Loop Is Required for Full Kinase Activity In Vivo of Human and Yeast Eukaryotic Initiation Factor 2α Kinases PKR and GCN2. Mol. Cell. Biol. 18, 2282–2297 (1998).

35. B. Nolen, S. Taylor, G. Ghosh, Regulation of protein kinases; controlling activity through activation segment conformation. Mol. Cell 15, 661–675 (2004).

36. R. Reinhardt, T. A. Leonard, A critical evaluation of protein kinase regulation by activation loop autophosphorylation. eLife 12, e88210.

37. T. Ito, A. Marintchev, G. Wagner, Solution structure of human initiation factor eIF2alpha reveals homology to the elongation factor eEF1B. Structure 12, 1693–1704 (2004).

38. A. C. Dar, T. E. Dever, F. Sicheri, Higher-order substrate recognition of eIF2alpha by the RNA-dependent protein kinase PKR. Cell 122, 887–900 (2005).

39. C. A. Dodson, S. Yeoh, T. Haq, R. Bayliss, A kinetic test characterizes kinase intramolecular and intermolecular autophosphorylation mechanisms. Sci. Signal. 6, ra54 (2013).

40. S. J. Marciniak, L. Garcia-Bonilla, J. Hu, H. P. Harding, D. Ron, Activation-dependent substrate recruitment by the eukaryotic translation initiation factor 2 kinase PERK. J. Cell Biol. 172, 201–209 (2006).

41. A. M. Darnell, A. R. Subramaniam, E. K. O’Shea, Translational Control through Differential Ribosome Pausing during Amino Acid Limitation in Mammalian Cells. Mol. Cell 71, 229–243.e11 (2018).

42. J. Misra, M. J. Holmes, E. T Mirek, M. Langevin, H.-G. Kim, K. R. Carlson, M. Watford, X. C. Dong, T. G. Anthony, R. C. Wek, Discordant regulation of eIF2 kinase GCN2 and mTORC1 during nutrient stress. Nucleic Acids Res. 49, 5726–5742 (2021).

43. K. Sakamoto, A. Hayashi, A. Sakamoto, D. Kiga, H. Nakayama, A. Soma, T. Kobayashi, M. Kitabatake, K. Takio, K. Saito, M. Shirouzu, I. Hirao, S. Yokoyama, Site-specific incorporation of an unnatural amino acid into proteins in mammalian cells. Nucleic Acids Res. 30, 4692–4699 (2002).

44. A. Chatterjee, J. Guo, H. S. Lee, P. G. Schultz, A genetically encoded fluorescent probe in mammalian cells. J. Am. Chem. Soc. 135, 12540–12543 (2013).

45. I. Coin, V. Katritch, T. Sun, Z. Xiang, F. Y. Siu, M. Beyermann, R. C. Stevens, L. Wang, Genetically encoded chemical probes in cells reveal the binding path of urocortin-I to CRF class B GPCR. Cell 155, 1258–1269 (2013).

46. M. Ramirez, R. C. Wek, A. G. Hinnebusch, Ribosome association of GCN2 protein kinase, a translational activator of the GCN4 gene of Saccharomyces cerevisiae. Mol. Cell. Biol. 11, 3027–3036 (1991).

47. J. Sharma, M. Du, E. Wong, V. Mutyam, Y. Li, J. Chen, J. Wangen, K. Thrasher, L. Fu, N. Peng, L. Tang, K. Liu, B. Mathew, R. J. Bostwick, C. E. Augelli-Szafran, H. Bihler, F. Liang, J. Mahiou, J. Saltz, A. Rab, J. Hong, E. J. Sorscher, E. M. Mendenhall, C. J. Coppola, K. M. Keeling, R. Green, M. Mense, M. J. Suto, S. M. Rowe, D. M. Bedwell, A small molecule that induces translational readthrough of CFTR nonsense mutations by eRF1 depletion. Nat. Commun. 12, 4358 (2021).

48. J. P. L. Coelho, M. C. J. Yip, K. Oltion, J. Taunton, S. Shao, The eRF1 degrader SRI-41315 acts as a molecular glue at the ribosomal decoding center. Nat. Chem. Biol. 20, 877–884 (2024).

49. H. Qiu, C. Hu, J. Dong, A. G. Hinnebusch, Mutations that bypass tRNA binding activate the intrinsically defective kinase domain in GCN2. Genes Dev. 16, 1271–1280 (2002).

50. M. Huse, J. Kuriyan, The Conformational Plasticity of Protein Kinases. Cell 109, 275–282 (2002).

51. M. Dey, C. Cao, A. C. Dar, T. Tamura, K. Ozato, F. Sicheri, T. E. Dever, Mechanistic link between PKR dimerization, autophosphorylation, and eIF2alpha substrate recognition. Cell 122, 901–913 (2005).

52. H. Shinohara, M. Behar, K. Inoue, M. Hiroshima, T. Yasuda, T. Nagashima, S. Kimura, H. Sanjo, S. Maeda, N. Yumoto, S. Ki, S. Akira, Y. Sako, A. Hoffmann, T. Kurosaki, M. Okada-Hatakeyama, Positive feedback within a kinase signaling complex functions as a switch mechanism for NF-κB activation. Science 344, 760–764 (2014).

53. H. Mellor, C. G. Proud, A synthetic peptide substrate for initiation factor-2 kinases. Biochem. Biophys. Res. Commun. 178, 430–437 (1991).

54. M. Otto, T. A. Leonard, Dimerization-dependent cis-autophosphorylation activates the UPR and ISR kinases. bioRxiv [Preprint] (2026). 10.64898/2026.07.28.741179.

55. R. R. Adams, S. P. Wheatley, A. M. Gouldsworthy, S. E. Kandels-Lewis, M. Carmena, C. Smythe, D. L. Gerloff, W. C. Earnshaw, INCENP binds the Aurora-related kinase AIRK2 and is required to target it to chromosomes, the central spindle and cleavage furrow. Curr. Biol. CB 10, 1075–1078 (2000).

56. J. D. Bishop, J. M. Schumacher, Phosphorylation of the carboxyl terminus of inner centromere protein (INCENP) by the Aurora B Kinase stimulates Aurora B kinase activity. J. Biol. Chem. 277, 27577–27580 (2002).

57. F. Sessa, M. Mapelli, C. Ciferri, C. Tarricone, L. B. Areces, T. R. Schneider, P. T. Stukenberg, A. Musacchio, Mechanism of Aurora B Activation by INCENP and Inhibition by Hesperadin. Mol. Cell 18, 379–391 (2005).

58. C. Y. Chen, S. B. Dion, C. M. Kim, J. L. Benovic, Beta-adrenergic receptor kinase. Agonist-dependent receptor binding promotes kinase activation. J. Biol. Chem. 268, 7825–7831 (1993).

59. Q. Chen, M. Plasencia, Z. Li, S. Mukherjee, D. Patra, C.-L. Chen, T. Klose, X.-Q. Yao, A. A. Kossiakoff, L. Chang, P. C. Andrews, J. J. G. Tesmer, Structures of rhodopsin in complex with G-protein-coupled receptor kinase 1. Nature 595, 600–605 (2021).

60. J. Liu, A. Sudom, X. Min, Z. Cao, X. Gao, M. Ayres, F. Lee, P. Cao, S. Johnstone, O. Plotnikova, N. Walker, G. Chen, Z. Wang, Structure of the Nuclear Factor κB-inducing Kinase (NIK) Kinase Domain Reveals a Constitutively Active Conformation. J. Biol. Chem. 287, 27326–27334 (2012).

61. G. de Leon-Boenig, K. K. Bowman, J. A. Feng, T. Crawford, C. Everett, Y. Franke, A. Oh, M. Stanley, S. T. Staben, M. A. Starovasnik, H. J. A. Wallweber, J. Wu, L. C. Wu, A. R. Johnson, S. G. Hymowitz, The crystal structure of the catalytic domain of the NF-κB inducing kinase reveals a narrow but flexible active site. Structure 20, 1704–1714 (2012).

62. D. T. Hughes, M. Halliday, H. L. Smith, N. C. Verity, C. Molloy, H. Radford, A. J. Butcher, G. R. Mallucci, Targeting the kinase insert loop of PERK selectively modulates PERK signaling without systemic toxicity in mice. Sci. Signal. 13, eabb4749 (2020).

63. K. K. Brar, D. T. Hughes, J. L. Morris, K. Subramanian, S. Krishna, F. Gao, L.-S. Rieder, S. Uhrig, J. Freeman, H. L. Smith, R. Jukes-Jones, E. Avezov, J. Nunnari, J. Prudent, A. J. Butcher, G. R. Mallucci, PERK-ATAD3A interaction provides a subcellular safe haven for protein synthesis during ER stress. Science 385, eadp7114 (2024).

64. M. E. Evans, W. C. Clark, G. Zheng, T. Pan, Determination of tRNA aminoacylation levels by high-throughput sequencing. Nucleic Acids Res. 45, e133 (2017).

65. A. Behrens, G. Rodschinka, D. D. Nedialkova, High-resolution quantitative profiling of tRNA abundance and modification status in eukaryotes by mim-tRNAseq. Mol. Cell 81, 1802–1815.e7 (2021).

66. H. Paternoga, L. Xia, L. Dimitrova-Paternoga, S. Li, L. L. Yan, M. Oestereich, S. Kasvandik, A. N. Nanjaraj Urs, B. Beckert, T. Tenson, H. Zaher, T. Inada, D. N. Wilson, Structure of a Gcn2 dimer in complex with the large 60S ribosomal subunit. Proc. Natl. Acad. Sci. 122, e2415807122 (2025).

67. L. L. Yan, H. S. Zaher, Ribosome quality control antagonizes the activation of the integrated stress response on colliding ribosomes. Mol. Cell 81, 614–628.e4 (2021).

68. L. D. Kapp, J. R. Lorsch, GTP-dependent Recognition of the Methionine Moiety on Initiator tRNA by Translation Factor eIF2. J. Mol. Biol. 335, 923–936 (2004).

69. F. L. Erickson, E. M. Hannig, Ligand interactions with eukaryotic translation initiation factor 2: role of the gamma-subunit. EMBO J. 15, 6311–6320 (1996).

70. M. Naveau, C. Lazennec-Schurdevin, M. Panvert, E. Dubiez, Y. Mechulam, E. Schmitt, Roles of yeast eIF2α and eIF2β subunits in the binding of the initiator methionyl-tRNA. Nucleic Acids Res. 41, 1047–1057 (2013).

71. A. Thakur, S. Gaikwad, A. K. Vijjamarri, A. G. Hinnebusch, eIF2α interactions with mRNA control accurate start codon selection by the translation preinitiation complex. Nucleic Acids Res. 48, 10280–10296 (2020).

72. M. Szaruga, D. A. Janssen, C. de Miguel, G. Hodgson, A. Fatalska, A. P. Pitera, A. Andreeva, A. Bertolotti, Activation of the integrated stress response by inhibitors of its kinases. Nat. Commun. 14, 5535 (2023).

73. K. R. Carlson, M. M. Georgiadis, F. Tameire, K. A. Staschke, R. C. Wek, Activation of Gcn2 by small molecules designed to be inhibitors. J. Biol. Chem. 299, 104595 (2023).

74. T. Ito, G. Wagner, Using codon optimization, chaperone co-expression, and rational mutagenesis for production and NMR assignments of human eIF2 alpha. J. Biomol. NMR 28, 357–367 (2004).

75. Z. Huang, F. F. Diehl, M. Wang, Y. Li, A. Song, F. X. Chen, N. A. Rosa-Mercado, R. Beckmann, R. Green, J. Cheng, RIOK3 mediates the degradation of 40S ribosomes. Mol. Cell 85, 802–814.e12 (2025).

76. A. Mandlbauer, Q. Sun, N. Popitsch, T. Schwickert, M. Spanova, J. Wang, S. L. Ameres, M. Busslinger, L. Cochella, Mime-seq 2.0: a method to sequence microRNAs from specific mouse cell types. EMBO J. 43, 2506–2525 (2024).

77. E. Damoc, C. S. Fraser, M. Zhou, H. Videler, G. L. Mayeur, J. W. B. Hershey, J. A. Doudna, C. V. Robinson, J. A. Leary, Structural Characterization of the Human Eukaryotic Initiation Factor 3 Protein Complex by Mass Spectrometry*. Mol. Cell. Proteomics 6, 1135–1146 (2007).

78. C. S. Fraser, K. E. Berry, J. W. B. Hershey, J. A. Doudna, eIF3j is located in the decoding center of the human 40S ribosomal subunit. Mol. Cell 26, 811–819 (2007).

79. J. Abramson, J. Adler, J. Dunger, R. Evans, T. Green, A. Pritzel, O. Ronneberger, L. Willmore, A. J. Ballard, J. Bambrick, S. W. Bodenstein, D. A. Evans, C.-C. Hung, M. O’Neill, D. Reiman, K. Tunyasuvunakool, Z. Wu, A. Žemgulytė, E. Arvaniti, C. Beattie, O. Bertolli, A. Bridgland, A. Cherepanov, M. Congreve, A. I. Cowen-Rivers, A. Cowie, M. Figurnov, F. B. Fuchs, H. Gladman, R. Jain, Y. A. Khan, C. M. R. Low, K. Perlin, A. Potapenko, P. Savy, S. Singh, A. Stecula, A. Thillaisundaram, C. Tong, S. Yakneen, E. D. Zhong, M. Zielinski, A. Žídek, V. Bapst, P. Kohli, M. Jaderberg, D. Hassabis, J. M. Jumper, Accurate structure prediction of biomolecular interactions with AlphaFold 3. Nature 630, 493–500 (2024).

80. F. Sievers, A. Wilm, D. Dineen, T. J. Gibson, K. Karplus, W. Li, R. Lopez, H. McWilliam, M. Remmert, J. Söding, J. D. Thompson, D. G. Higgins, Fast, scalable generation of high-quality protein multiple sequence alignments using Clustal Omega. Mol. Syst. Biol. 7, 539 (2011).

81. Q. Tian, C. Wang, Y. Liu, W. Xie, Structural basis for recognition of G-1-containing tRNA by histidyl-tRNA synthetase. Nucleic Acids Res. 43, 2980–2990 (2015).

